# JMJD3 mediated senescence is required to overcome stress induced hematopoietic defects

**DOI:** 10.1101/2024.08.08.607138

**Authors:** Yuichiro Nakata, Takeshi Ueda, Yasuyuki Sera, Akinori Kanai, Ken-ichiro Ikeda, Norimasa Yamasaki, Akiko Nagamachi, Kohei Kobatake, Masataka Taguchi, Yusuke Sotomaru, Tatsuo Ichinohe, Zen-ichiro Honda, Ichiro Manabe, Toshio Suda, Keiyo Takubo, Osamu Kaminuma, Hiroaki Honda

## Abstract

Cellular senescence in stem cells compromises regenerative capacity, promotes chronic inflammation, and is implicated in aging. Hematopoietic stem and progenitor cells (HSPCs) are responsible for producing mature blood cells, however, how cellular senescence influences their function is largely unknown. Here, we show that JMJD3, a histone demethylase, activates cellular senescence via *p16^Ink4a^* upregulation in competition with Polycomb group proteins, and reprograms HSPC integrity to overcome hematopoietic defects induced by replicative and oncogenic stresses. JMJD3 deficiency impaired stem cell potential, proper cell cycle regulation, and WNT pathway activation in HSPCs under stress conditions. These impaired phenotypes were rescued through exogenous and retroviral introduction of *p16^Ink4a^*. This JMJD3-p16^INK4a^ axis in hematopoiesis is age-dependent and is distinct from cellular senescence. Treatment with a JMJD3 selective inhibitor attenuated leukemic potential during cellular senescence. Taken together, these results demonstrate that JMJD3-p16^INK4a^ mediates cellular senescence and plays critical roles in the functional integrity of HSPCs under stress conditions, suggesting a new link for aging and anti-cancer therapies.

## Introduction

Hematopoietic stem and progenitor cells (HSPCs) are capable of self-renewal and differentiation for all blood cell lineages (Orkin *et al*, 2008). Maintaining HSPC stem cell potential is required for supplying mature blood cells throughout the lifetime of individual. Regulatory machineries to balance proliferative and non-proliferative status in HSPCs are critical to avoid exhaustion and to sustain stem cell pools (Li, 2011). Stable cell cycle arrest without stimuli, called as quiescence, is mediated by specific cell cycle regulators, such as p27^KIP1^ and p57^KIP2^, are well characterized in HSPCs (Zou *et al*, 2011; Matsumoto *et al*, 2011). Cellular senescence is considered to be a type of cell cycle arrest in response to various intrinsic and extrinsic stimuli and is associated with attenuation of stem cell capacity, release of inflammatory cytokines, and spread of senescent cells (Huang *et al*, 2022). On the other hand, cellular senescence also promotes biological benefits including enhanced kidney regenerative capacity (DiRocco *et al*, 2014) and promoting insulin production in pancreatic beta cells (Helman *et al*, 2016). These reports suggest that cellular senescence mediates stem cell fate determination though intricate transcriptional reprogramming and may be tissue- or cell-type dependent. As such, cellular senescence may control stem cell activity in hematopoiesis, though the mechanisms remain unclear.

*CDKN2A*, encoding p16^INK4a^ and p19^ARF^, is a key senescence gene. P16^INK4a^ negatively regulates cell cycle progression by binding to CDK4/CDK6, thereby inhibiting CyclinD1-CDK4/CDK6 complex formation which phosphorylates RB resulting in release of the cell cycle promoter, E2F (Ewen *et al*, 1994). The *CDKN2A locus* is epigenetically regulated by Polycomb group proteins, including Polycomb repressive complex (PRC) 1 and 2 (Neff *et al*, 2012; Mohammad *et al*, 2017; Biehs *et al*, 2013; Bruggeman *et al*, 2007). PRC2, which consists of catalytic (EZH2) and non-catalytic (EED and SUZ12) subunits, recognizes trimethylated lysine 27 on histone H3 (H3K27me3) and contributes to gene silencing (Simon *et al*, 2009). Cell fate of HSPCs is controlled by transcription factors in combination with epigenetic regulators, such as PRC2. Previous reports showed that PRC2 plays essential roles in the functional integrity of HSPCs by regulating expression of target genes (Di Carlo *et al*, 2019; Margueron *et al*, 2011; Radulović *et al*, 2013). Thus, perturbation of PRC2 function, along with compromised H3K27me3, is directly linked to loss of HSPC activity and/or leukemogenesis. In fact, gain- and loss-of-function mutations in PRC2 constituents were identified in various hematopoietic malignancies (Lohr *et al*, 2012; Nikoloski *et al*, 2010; Ueda *et al*, 2012).

Two distinct demethylases for H3K27 were identified, UTX/KDM6A and JMJD3/KDM6B. These enzymes share a JmjC domain that catalyzes histone lysine demethylation (Hong *et al*, 2007; Klose *et al*, 2006). UTX functions as a tumor suppressor in various cancers including hematopoietic malignancies (Mar *et al*, 2012; van Haaften *et al*, 2009). In contrast, JMJD3 is highly expressed in hematopoietic disorders (Ntziachristos *et al*, 2014; Ohguchi *et al*, 2017; Wei *et al*, 2013). Accordingly, these two enzymes may have distinct functions that target different genes in adult hematopoiesis. Indeed, a previous report demonstrated that JMJD3 and UTX play contrasting roles in acute lymphoblastic leukemia (Ntziachristos *et al*, 2014). JMJD3 was discovered as a key regulator of macrophages under inflammatory and differentiation stimuli (De Santa *et al*, 2007). Subsequent reports demonstrated that JMJD3 regulates inflammatory gene loci and macrophage polarization through its demethylase activity (De Santa *et al*, 2009; Satoh *et al*, 2010). In addition, JMJD3 plays crucial roles in the differentiation and maintenance of various types of stem cells such as embryonic stem cells, mesenchymal stem cells, and neural stem cells (Ohtani *et al*, 2013; Park *et al*, 2014; Ye *et al*, 2012), and its ectopic expression accelerates the differentiation of human induced pluripotent stem cells (iPSCs) (Akiyama *et al*, 2016).

JMJD3 is also recruited to the *CDKN2A locus* and induces cellular senescence to prevent cancer cell proliferation in response to stress (Lin *et al*, 2012; He *et al*, 2017; Agger *et al*, 2009; Barradas *et al*, 2009). Given that the JMJD3-p16^INK4a^ axis is essential for cellular senescence, we expect that Polycomb group proteins and JMJD3 competitively fine-tune expression of *p16^INK4a^* via methylation and demethylation activities to control cellular senescence in HSPCs. However, whether the JMJD3-p16^INK4a^ axis induces cellular senescence in hematopoiesis and how it influences HSPC potential are still mostly unknown. In this study, we generated conditional *Jmjd3* knockout mice and demonstrated that JMJD3 epigenetically regulates expression of *p16^INK4a^* under stress conditions as a senescence inducer and is associated with maintenance of HSPC integrity by gene reprogramming and protection against excessive cell cycle entry.

## Results

### Acquired deletion of JMJD3 induces minimal defects on adult hematopoiesis at steady state

First, to investigate the role of JMJD3 in steady state hematopoiesis, we generated mice in which exons 15–17 of the *Jmjd3* gene, which encode part of the JmjC domain, were flanked by two *loxP* sites (Fig EV1A and EV1B). pIpC treatment almost completely deleted exons 15–17 derived transcripts and JMJD3 protein in *Jmjd3^flox/flox^*;*MxCre*^+^ BM cells (Fig EV1C), indicating successful ablation of the *Jmjd3* gene product in the hematopoietic system (hereafter, pIpC-treated *Jmjd3^flox/flox^;MxCre*^−^ and *Jmjd3^flox/flox^;MxCre*^+^ mice are referred to as *Jmjd3^+/+^* and *Jmjd3^Δ/Δ^* mice, respectively).

Analysis of the PB parameters of *Jmjd3^+/+^* and *Jmjd3^Δ/Δ^* mice showed no obvious changes in white blood cell (WBC) counts, hemoglobin (Hgb) concentration, platelet (Plt) number, or differentiation status of WBCs (Fig EV2A and EV2B). In addition, percentage of the lineage negative (Lin^−^) population and cell numbers of HSPC subfractions (Table EV1) in the BM were similar between the two groups, except for a slight decrease in CMP and MEP fractions in *Jmjd3^Δ/Δ^* mice (Fig EV2C and EV2D). No hematological diseases developed in *Jmjd3^Δ/Δ^* mice during the 1.5 year observation period. These results indicate that *Jmjd3* deficiency does not induce obvious changes in steady state hematopoiesis.

### Loss of JMJD3 impairs long-term repopulating activity of HSPCs under BMT induced stress conditions

Since no apparent changes were observed under steady state hematopoiesis, we next examined the behavior of *Jmjd3^Δ/Δ^*HSPCs under stress conditions. We first performed serial bone marrow transplantation (BMT) experiments to assess the role of JMJD3 under replicative stress (Fig 1A). Equal numbers of Lin^−^, Sca-1^+^ and c-kit^+^ (LSK) cells from *Jmjd3^+/+^* and *Jmjd3^Δ/Δ^* mice, which contain similar number of HSCs (Fig EV2D), were transplanted into lethally irradiated syngeneic mice. In the 1^st^ transplant, PB chimerism, PB differentiation status, and percentage of the Lin^−^ population in donor derived BM cells were similar in recipients transplanted with *Jmjd3^+/+^* and *Jmjd3^Δ/Δ^* LSK cells, although donor-derived chimerism in various BM subfractions was higher in mice transplanted with *Jmjd3^Δ/Δ^* cells (Fig 1B). In contrast, *Jmjd3^Δ/Δ^* cells of the 2^nd^ transplant exhibited significantly lower PB chimerism without affecting differentiation, significantly lower chimerism in all BM subfractions, and a markedly reduced Lin^−^ population in donor-derived BM cells (Fig 1C).

**Figure 1.**
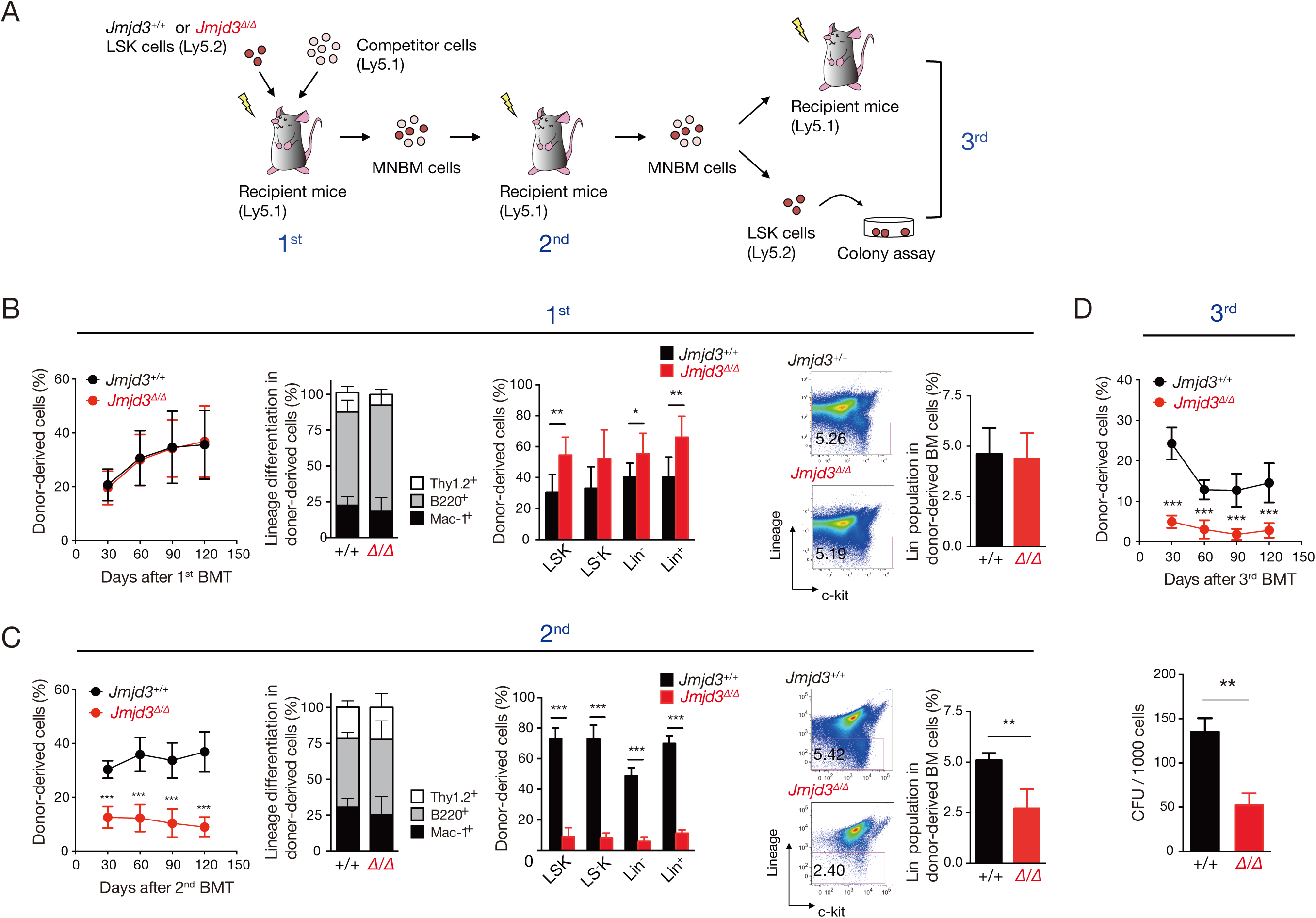
Analysis of *Jmjd3^Δ/Δ^* HSPCs in replicative stress. (A) Schematic diagram of the serial competitive BM reconstitution experiment. 2.0×10^3^ LSK cells from *Jmjd3^+/+^*and *Jmjd3^Δ/Δ^* mice were transplanted into lethally irradiated primary recipients with 2.5×10^5^ wildtype competitor mononuclear BM (MNBM) cells. 3.0×10^6^ MNBM cells from the first BMT recipients were transplanted in lethally irradiated secondary recipients, and 3.0×10^6^ MNBM cells from second recipients were subjected to transplantation into tertiary recipients and colony formation assays. (B) Chimerism and lineage differentiation of donor-derived cells in the PB of the 1^st^ recipients (left and middle left panels, n = 10 each) and chimerism of donor-derived cells in BM subfractions including LSK, LS^−^K (Lin^−^, Sca-1^−^, c-kit^+^), Lin^−^, and Lin^+^ of 1^st^ recipients at 4 months after BMT (middle right panel) (mean ± SD, n = 6). Flow cytometric profiles of donor-derived Lin^−^ cells in the BM of the 1^st^ BMT recipients 4 months after BMT (right panel) (mean + SD, n = 5). (C) The same analyses in the 2^nd^ recipients (mean ± SD, n = 12). (D) Chimerism of donor-derived cells in the PB of 3^rd^ recipients (mean ± SD, n = 11) and colony formation assays of LSK cells from the 3^rd^ recipients at 4 months (mean + SD, n = 3). \**p* < 0.05, \*\**p* < 0.01, \*\*\**p* < 0.001.

To further investigate the long-term effects of JMJD3 deletion on HSPC activity, donor-derived BM cells from the second transplant were subjected to a 3^rd^ transplant and colony formation assays were conducted from LSK cells. A significant reduction of donor-derived PB chimerism was observed in 3^rd^ transplants with *Jmjd3^Δ/Δ^* cells, and *Jmjd3^Δ/Δ^*LSK cells generated significantly fewer colonies (Fig 1D). These results indicate that loss of JMJD3 transiently confers a proliferative activity on HSPCs but eventually impairs their long-term repopulating and colony-forming abilities.

### Loss of JMJD3 impairs leukemogenic activity of HSPCs under oncogene induced stress conditions

We next examined the role of JMJD3 under oncogenic stress. We introduced *MLL-AF9*, a well-known leukemogenic gene that induces acute myeloid leukemia (AML), coupled with *EGFP* into Lin^−^, c-kit^+^ (LK) cells, and EGFP^+^ cells were subjected to colony replating and transplantation assays (Fig 2A). At the second round of colony replating, *MLL-AF9* expressing (*MA9*) *Jmjd3^Δ/Δ^* cells generated slightly more colonies but by the third round of replating, *MA9 Jmjd3^Δ/Δ^* colony numbers were significantly reduced compared with *MA9 Jmjd3^+/+^* (Fig 2B). In addition, the percentage of LK cells in *MA9 Jmjd3^Δ/Δ^* colonies decreased with replating (Fig 2C). To evaluate *in vivo* tumorigenic activity, *MA9 Jmjd3^+/+^* and *Jmjd3^Δ/Δ^* cells were transplanted into lethally irradiated syngeneic mice. Significantly longer survival was observed in recipients transplanted with *MA9 Jmjd3^Δ/Δ^* cells compared with *MA9 Jmjd3^+/+^* cells (Fig 2D), as well as a significantly lower percentage of Lin^−^ cells in the BM (Fig 2E).

**Figure 2.**
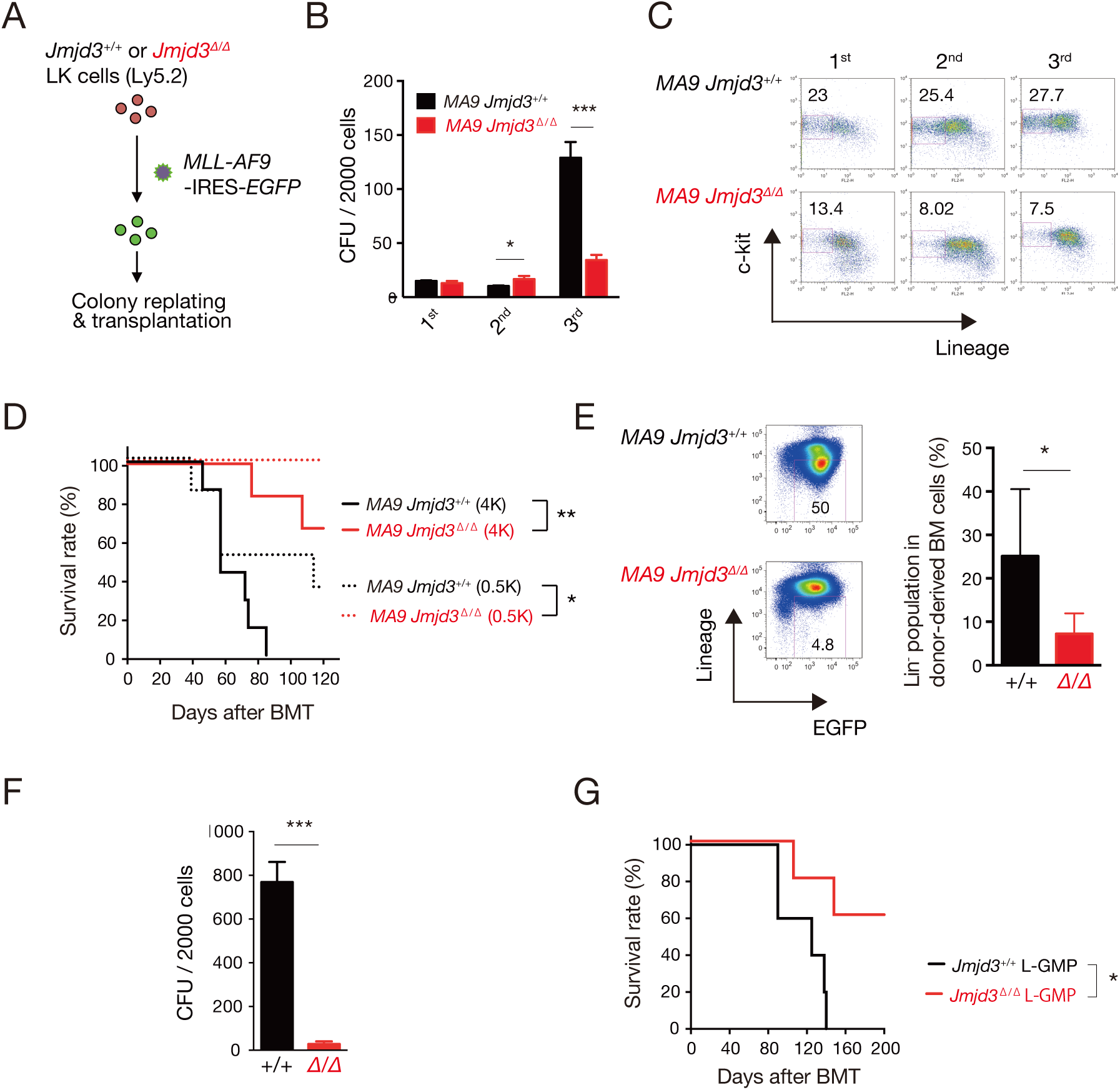
Analysis of *Jmjd3^Δ/Δ^* HSPCs in oncogenic stress. (A) Schematic diagram of *MLL-AF9* oncogene transduction. LK cells from *Jmjd3^+/+^* and *Jmjd3^Δ/Δ^* mice were transfected with *MLL-AF9-*IRES*-EGFP* retrovirus, and EGFP^+^ cells were subjected to the following assays. (B) Colony replating assays. Bars indicate colony number (CFU; colony-forming unit) in each round of plating (mean + SD, n = 3). (C) Flow cytometric profiles of colony-forming cells. Cells were stained with c-kit and lineage markers, and the percentages of c-kit^+^, Lin^−^ cells in each round are shown. (D) Kaplan–Meier survival curves of mice transplanted with *MLL-AF9* expressing *Jmjd3^+/+^* and *Jmjd3^Δ/Δ^* cells. 4.0×10^3^ (4K) or 5.0×10^2^ (0.5K) cells were transplanted into lethally irradiated recipients with 2.5×10^5^ wild type competitor MNBM cells (n = 6–7). (E) BM cells from mice that developed *MLL-AF9* induced leukemia were stained with EGFP and lineage markers. The percentage of EGFP^+^, Lin^−^ cells is shown (mean + SD, n = 4). (F) Colony replating assay of *Jmjd3^+/+^* and *Jmjd3^Δ/Δ^* L-GMPs at the 3^rd^ round of replating (mean + SD, n = 3). (G) Kaplan–Meier survival curves of mice transplanted with *Jmjd3^+/+^* and *Jmjd3^Δ/Δ^* L-GMPs. 2.0×10^2^ L-GMPs were transplanted into lethally irradiated recipients with 2.5×10^5^ wild type competitor MNBM cells for radioprotection (n = 5). \**p* < 0.05, \*\**p* < 0.01, \*\*\**p* < 0.001.

Leukemia arises from leukemic stem cells (LSCs) and recent studies demonstrated that LSCs transformed by *MLL-AF9* were enriched in the GMP fraction, known as leukemic-GMP (L-GMP) (Krivtsov *et al*, 2006). Thus, we performed the same analyses using L-GMPs. Colony numbers of *Jmjd3^Δ/Δ^* L-GMPs were significantly lower than those of *Jmjd3^+/+^* L-GMPs (Fig 2F). In addition, recipients transplanted with *Jmjd3^Δ/Δ^* L-GMPs exhibited significantly lower mortality than those from *Jmjd3^+/+^* L-GMPs (Fig 2G). These findings collectively indicate that JMJD3 deficiency impairs *MLL-AF9* induced leukemogenesis by perturbing the leukemogenic potential of LSCs.

### JMJD3 epigenetically controls expression of *p16^Ink4a^* in HSPCs under stress conditions while competing with PRC2

Since *Cdkn2a* are known common targets of JMJD3 and Polycomb group proteins (Neff *et al*, 2012; Mohammad *et al*, 2017; Biehs *et al*, 2013; Bruggeman *et al*, 2007; Lin *et al*, 2012), we rationalized that p16^INK4a^ played a role in the hematopoietic deficiencies observed under stress conditions caused by loss of JMJD3. To investigate the molecular mechanisms underlying the impaired stem cell potential of *Jmjd3^Δ/Δ^* HSPCs under stress conditions and the relation with p16^INK4a^, we first examined expression levels of *CDKI* genes including *Cdkn2a* in three different types of cells including 1) LSK (Steady), LSK cells at steady state, 2) LSK (BMT), LSK cells after second BMT, and 3) L-GMP, GMP fraction from LK cells after *MLL-AF9* transduction from wild type mice. Drastic upregulation of *Cdkn2a* expression was observed in HSPCs under stress compared with those at steady state, suggesting that *Cdkn2a* has essential roles in hematopoiesis under stress (Fig 3A). Next, to investigate whether JMJD3 deficiency influences the upregulation of *p16^INK4a^*, we examined *CDKI* expression profiles in *Jmjd3^Δ/Δ^* cells. We found that expression levels of *p16^Ink4a^* and *p19^Arf^*, were markedly lower in *Jmjd3^Δ/Δ^* LSK (BMT) and L-GMP compared with *Jmjd3^+/+^* but not in *Jmjd3^Δ/Δ^* LSK (Steady) (Fig 3B).

**Figure 3.**
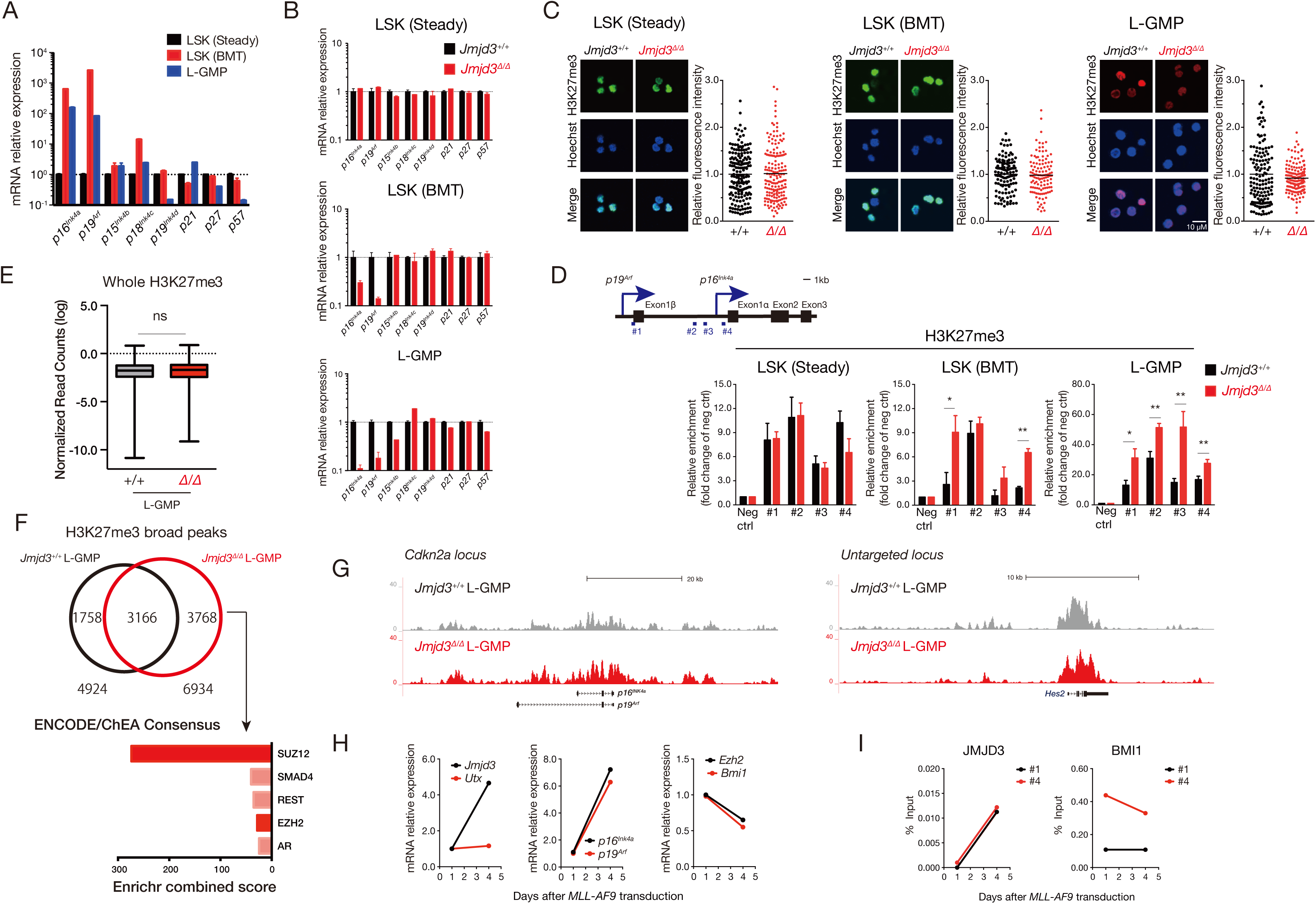
Demethylase-dependent regulation at the *Cdkn2a* locus in *Jmjd3^+/+^* and *Jmjd3^Δ/Δ^* HPSCs. (A) qPCR of *CDKI* genes in LSK (Steady), LSK (BMT), and L-GMP. Relative fold-changes to LSK (Steady) are shown on a logarithmic scale (mean + SD, n = 3). (B) qPCR of *CDKI* genes in LSK (Steady), LSK (BMT), and L-GMP of *Jmjd3^+/+^* and *Jmjd3^Δ/Δ^* mice (mean + SD, n = 3). (C) Immunofluorescence staining (left panels) and relative fluorescence intensity (right panels) of H3K27me3 in LSK (Steady), LSK (BMT), and L-GMP from *Jmjd3^+/+^* and *Jmjd3^Δ/Δ^*mice. Mean values are indicated as bars (n = 117-175). (D) Schematic diagram of the *Cdkn2a* locus. Promoter regions of *p19^Arf^* and *p16^Ink4a^* are indicated #1-4. (upper panel). H3K27me3 levels in the promoter regions of *p19^Arf^* and *p16^Ink4a^* genes in LSK (Steady), LSK (BMT), and L-GMP of *Jmjd3^+/+^* and *Jmjd3^Δ/Δ^* mice. Results are shown as fold changes relative to the negative control (Neg ctrl), (mean + SD, n = 3) (lower panel). (E) Box plot showing the global levels of H3K27me3 in L-GMPs from *Jmjd3^+/+^* and *Jmjd3^Δ/Δ^* mice. (F) Venn diagram showing the distributed broad H3K27me3 peaks in L-GMPs from *Jmjd3^+/+^*and *Jmjd3^Δ/Δ^* mice (upper panel) and enrichment analysis from unique H3K27me3 peaks *Jmjd3^Δ/Δ^* L-GMPs with Enrichr on the ENCODE/ChEA database (lower). (G) Genome browser view of H3K27me3 enrichment at *Cdkn2a* and an untargeted locus in L-GMPs from *Jmjd3^+/+^*and *Jmjd3^Δ/Δ^* mice. (H) qPCR analysis of H3K27 histone demethylases (*Jmjd3* and *Utx*) (left panel), *Cdkn2a* (*p16^Ink4a^*and *p19^Arf^*) (middle panel), and Polycomb genes (*Ezh2* and *Bmi1*) (right panel) in LK cells transduced with *MLL-AF9*. Expression levels are shown relative to day 1 (mean, n = 3). (I) ChIP-qPCR analysis for the enrichment of JMJD3 and BMI1 at the indicated *Cdkn2a* promoter regions in LK cells transduced with *MLL-AF9* (mean, n = 3) (#1 and #4, see Fig 3D).

Given that JMJD3 upregulates *p16^Ink4a^* expression through its demethylase activity on H3K27 (He *et al*, 2017; Agger *et al*, 2009; Barradas *et al*, 2009), we next investigated whether the suppression of *Cdkn2a* in stressed *Jmjd3^Δ/Δ^* HSPCs is linked to changes in H3K27me3. No obvious changes on global H3K27me3 levels were observed among LSK (Steady), LSK (BMT), and L-GMP from *Jmjd3^+/+^*and *Jmjd3^Δ/Δ^* mice by immunostaining (Fig 3C), prompting us to compare H3K27me3 enrichment in the promoter regions of *Cdkn2a* locus by ChIP-qPCR. No changes were observed in LSK (Steady), but significant enrichment of H3K27me3 was detected in the promoter regions of both *p16^Ink4a^* and *p19^Arf^* in *Jmjd3^Δ/Δ^* LSK (BMT) and L-GMP (Fig 3D). These data indicate that *Jmjd3* deficiency leads to insufficient demethylation of H3K27me3 at the *Cdkn2a* promoter in HSPCs under stress, which impairs expression of *p16^Ink4a^* and *p19^Arf^*, despite comparable global H3K27me3 levels. To further investigate the genome-wide distribution of H3K27me3, we performed Cut&Run sequencing in L-GMP from *Jmjd3^+/+^*and *Jmjd3^Δ/Δ^* mice. As expected, no obvious changes were observed on genome-wide H3K27me3 accumulation, however, we identified 3768 unique H3K27me3 peaks in *Jmjd3^Δ/Δ^* L-GMP. The genes annotated from these peaks were associated with PRC2 (SUZ12 and EZH2) target genes, indicating that JMJD3 and Polycomb group proteins may have common targetability and competitiveness in hematopoiesis (Fig 3E-G).

Our results suggest that JMJD3 may not only control expression of *Cdkn2a* genes but may also counteract H3K27me3 at the *Cdkn2a* locus mediated by Polycomb group proteins during stress conditions. To verify this, we examined expression changes of H3K27 demethylases (*Jmjd3* and *Utx*), *Cdkn2a* genes (*p16^Ink4a^* and *p19^Arf^*), and Polycomb group genes (*Ezh2* and *Bmi1*) following *MLL-AF9* transduction. *p16^Ink4a^*, *p19^Arf^,* and *Jmjd3*, but not *Utx*, were upregulated whereas *Ezh2* and *Bmi1* were downregulated (Fig 3H). In addition, ChIP-qPCR analysis revealed that upon *MLL-AF9* transduction, JMJD3 was recruited to the promoter regions of the *Cdkn2a* locus but BMI1 recruitment decreased (Fig 3I). These data indicate that JMJD3 and Polycomb group proteins control the expression of *Cdkn2a* genes competitively under stress conditions.

### Activation of the JMJD3-p16^INK4a^ axis protects HSPCs from cell cycle over-entry by inducing senescence

To investigate the JMJD3-p16^INK4a^ axis induced cellular phenotype on senescence and cell cycle progression, β-galactosidase staining assay, a marker of cellular senescence, was performed in LSK cells (Steady) and L-GMPs from *Jmjd3^+/+^*and *Jmjd3^Δ/Δ^* in combination with adriamycin (ADR) treatment to enhance the senescence induction. Although the population percentage of β-galactosidase^+^ cells between *Jmjd3^+/+^*and *Jmjd3^Δ/Δ^* LSK cells (Steady) were comparable, the ratio in *Jmjd3^Δ/Δ^* L-GMPs was significantly reduced (Fig 4A). Moreover, the percentage of β-galactosidase^+^ cells in L-GMP is much higher than in LSK (Steady) cells, strongly suggesting that cellular senescence in hematopoiesis may be essential for maintaining stem cell potential in response to stress but not at steady state. Next, we performed BrdU incorporation assays to investigate cell cycle progression. HSPCs under stress conditions with an impaired JMJD3-p16^INK4a^ axis exhibited significantly increased BrdU uptake compared with HSPCs at steady state (Fig 4B), indicating that the cellular senescence induced by activation of the JMJD3-p16^INK4a^ axis is crucial for maintaining stem cell activity in HSPCs under stress by regulating cell cycle entry.

**Figure 4.**
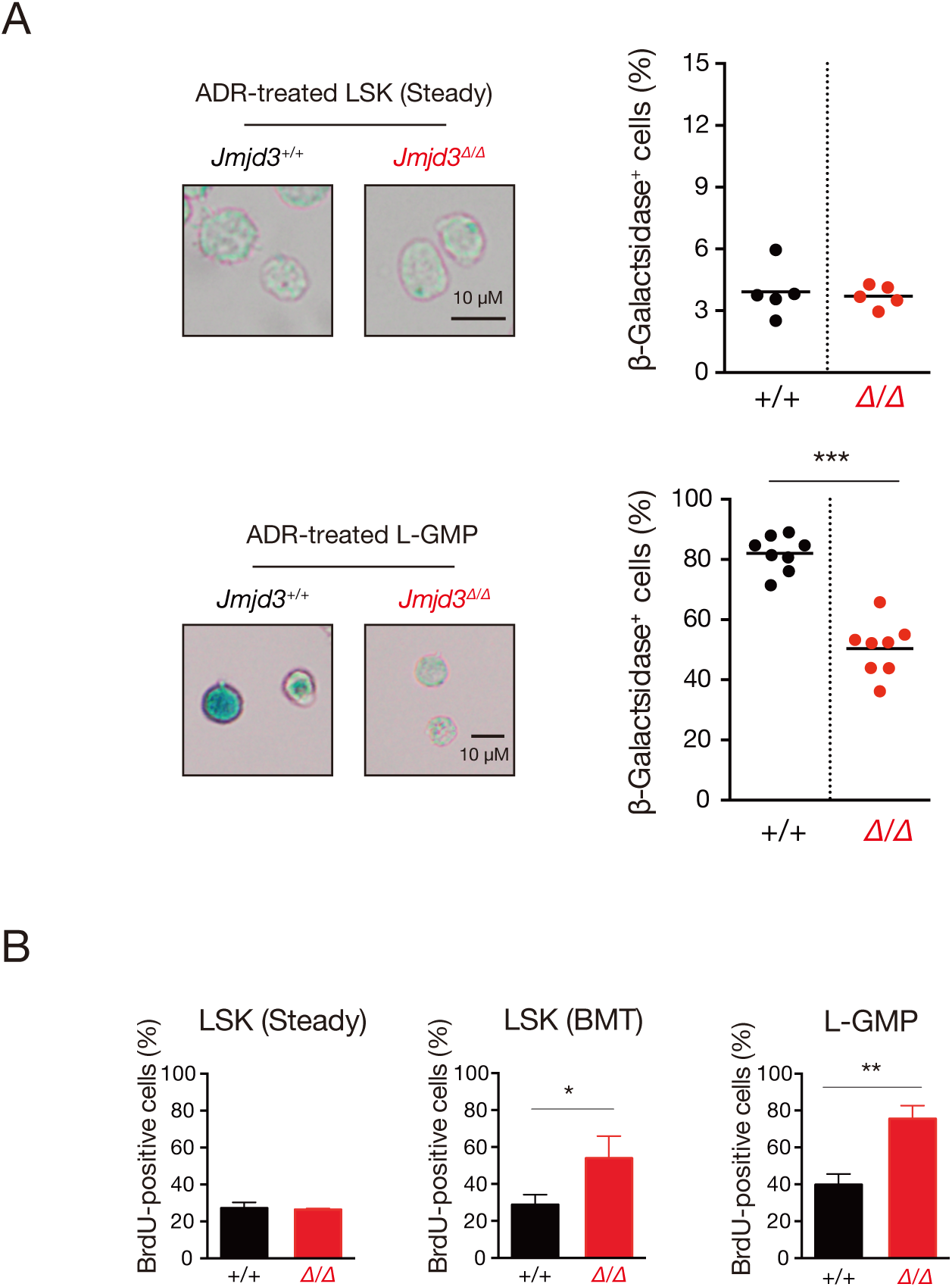
Jmjd3 deficiency failed induction of cellular senescence under stress condition. (A) *β*-galactosidase staining (left panel) and percentage of β-galactosidase^+^ cells in *Jmjd3^+/+^* and *Jmjd3^Δ/Δ^* LSK (Steady) (n = 4) and L-GMPs (n = 8) treated with adriamycin (ADR) (right panel). Mean values are indicated as bars. (B) Flow cytometric analysis of BrdU incorporation in LSK (Steady), LSK (BMT), and L-GMP of *Jmjd3^+/+^* and *Jmjd3^Δ/Δ^* mice (n = 3-4) (mean + SD). \**p* < 0.05, \*\**p* < 0.01, \*\*\**p* < 0.001.

### The JMJD3-p16^INK4a^ axis activates senescence induced reprogramming in HSPCs under stress conditions

To address global gene expression changes induced by JMJD3-p16^INK4a^ axis mediated senescence between *Jmjd3^+/+^* and *Jmjd3^Δ/Δ^* LSK (Steady), LSK (BMT), and L-GMP, we performed RNA-sequencing (RNA-seq). Genes with more than 2-fold up- or downregulation are shown in Fig 5A. Except for downregulated genes in LSK (BMT), expression levels in fewer than 2% of all genes were affected by *Jmjd3* deficiency, suggesting that although the defects of *Jmjd3^Δ/Δ^* HSPCs could be attributed to specific and limited JMJD3 target genes, *Jmjd3* deficiency under stress conditions greatly influences global gene expression changes compared with *Jmjd3* deficiency at steady state.

**Figure 5.**
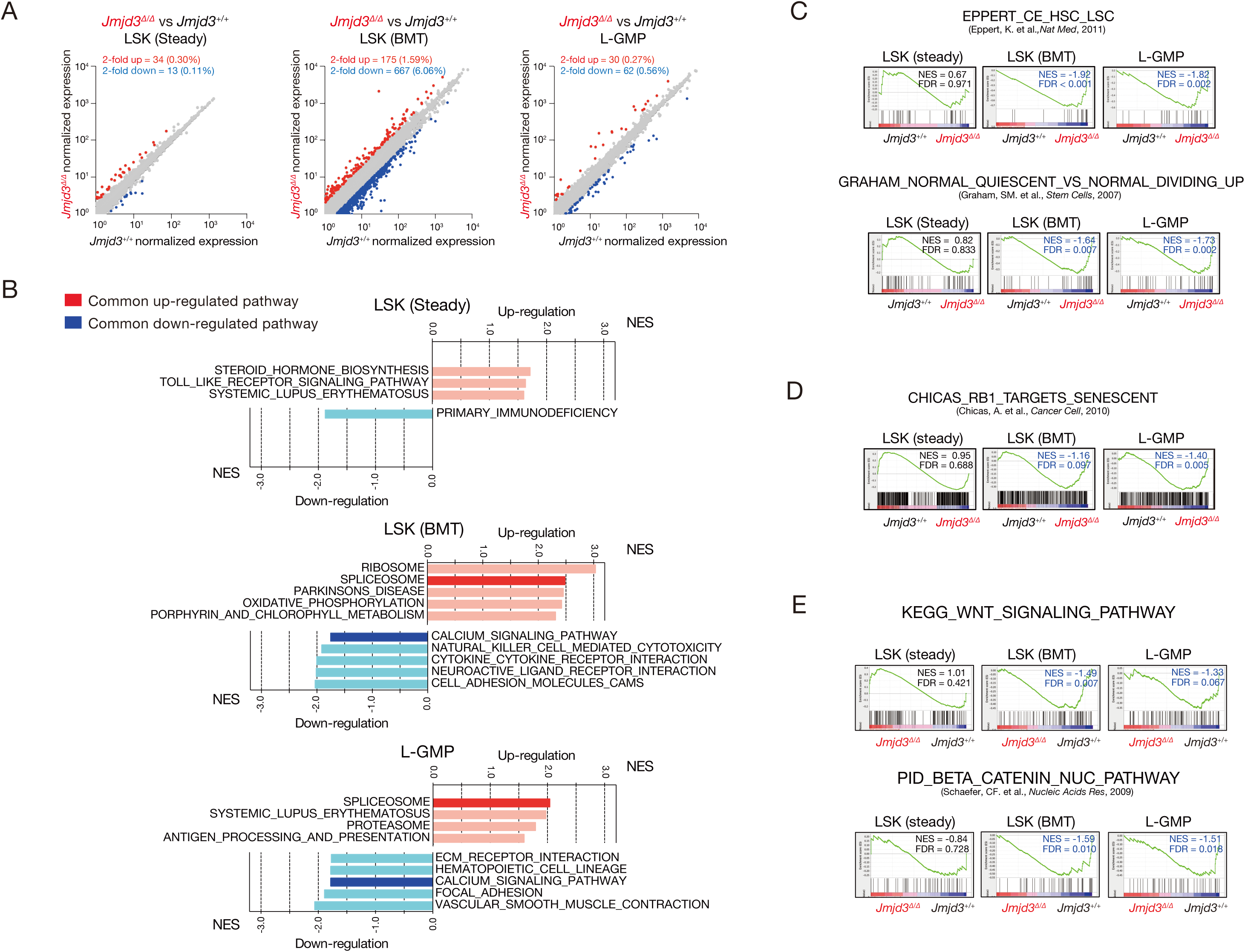
Expression and pathway analysis of senescence associated genes. (A) Scatter plots comparing normalized expression of individual genes (RPKM > 1) in LSK (Steady), LSK (BMT), and L-GMP of *Jmjd3^+/+^* and *Jmjd3^Δ/Δ^* mice. Numbers and percent of genes up- or downregulated more than 2-fold in *Jmjd3^Δ/Δ^* cells compared with *Jmjd3^+/+^*cells are shown as red and blue dots, respectively. (B) The top five most positively and negatively enriched KEGG pathways in LSK (Steady) (upper), LSK (BMT) (middle), and L-GMP (lower) from *Jmjd3^Δ/Δ^* mice compared with *Jmjd3^+/+^* mice (FDR < 0.25). Common up- and downregulated pathways between LSK (BMT) and L-GMP are shown as red and blue bars, respectively. (C) GSEA plots of LSK (Steady), LSK (BMT), and L-GMP in the indicated gene sets (top, genes commonly upregulated in human HSCs and LSCs; bottom, genes commonly upregulated in quiescent human CD34^+^ hematopoietic cells). Enrichment in *Jmjd3^Δ/Δ^*cells relative to *Jmjd3^+/+^* are shown with NES and FDR values. (D) GSEA plots of LSK (Steady), LSK (BMT), and L-GMP in genes commonly upregulated through a p16^INK4a^/RB1 pathway. Results are shown with NES and FDR values. (E) GSEA plots of WNT related pathways (KEGG_WNT_SIGNALING_PATHWAY (upper) and PID_BETA_CATENIN_NUC_PATHWAY (lower) in LSK (Steady), LSK (BMT), and L-GMP. *Jmjd3^Δ/Δ^* cells compared with *Jmjd3^+/+^* are shown with NES and FDR values.

We then assessed pathway changes using Gene Set Enrichment Analysis (GSEA). Among KEGG gene sets, we identified both up- and downregulated biological pathways (Fig 5B). Upregulation of SPLICEOSOME and downregulation of CALCIUM_SIGNALING_PATHWAY gene sets were commonly observed in *Jmjd3^Δ/Δ^*LSK (BMT) and L-GMP (indicated by red and blue bars, respectively), suggesting that these pathways are affected by *Jmjd3* deficiency under stress conditions. Notably, both *Jmjd3^Δ/Δ^* LSK (BMT) and L-GMP showed significant negative enrichment of genes that are upregulated in HSCs and LSCs (EPPERT_CE_HSC_LSC) (Eppert *et al*, 2011), as well as genes upregulated in quiescent HSCs (GRAHAM_NORMAL_QUIESCENT_VS_NORMAL_DIVIDING_UP) (Graham *et al*, 2007). These expression changes were not observed in *Jmjd3^Δ/Δ^* LSK (Steady) (Fig 5C). Consistent with Figures 1, 2 and 4, these findings indicate that *Jmjd3* deficiency impairs stem cell features by perturbing quiescence of HSCs and LSCs and inducing excessive HSPC and LSC proliferation under stress. To emphasize that JMJD3 and Polycomb group proteins competitively regulate their target loci by via enzymatic activities under stress conditions, we investigated possible target gene overlaps between JMJD3 and Polycomb proteins (Bracken *et al*, 2006; Wiederschain *et al*, 2007; Douglas *et al*, 2008; Ben-Porath *et al*, 2008; Kondo *et al*, 2008). We found that several gene sets upregulated in Polycomb deficient cells were significantly negatively enriched in *Jmjd3^Δ/Δ^*LSK (BMT) and L-GMP (Fig EV3), indicating that this substantial overlap in JMJD3 and Polycomb target genes may enable cells to fine-tune gene expression quickly and reversibly when stress is induced.

As expected, the p16^INK4a^/RB1 senescent pathway (CHICAS_RB1_TARGET_SENESCENT) (Chicas *et al*, 2010) was negatively enriched in *Jmjd3^Δ/Δ^* LSK (BMT) and L-GMP, but not in *Jmjd3^Δ/Δ^* LSK (Steady) (Fig 5D). A major characteristic of senescence associated reprogramming is significant enrichment of canonical WNT signaling (Milanovic *et al*, 2018). WNT SIGNALING and BETA CATENIN NUC (Schaefer *et al*, 2008) were negatively enriched in stress induced *Jmjd3^Δ/Δ^* HSPCs (LSK (BMT) and L-GMP) (Fig 5E). These findings strongly suggest that the JMJD3-p16^INK4a^ axis is a critical regulator of senescence associated reprogramming and contributes to maintaining the stemness of HSPCs under stress conditions.

### Exogenous p16^INK4a^ rescues impaired repopulating potential of *Jmjd3* deficient HSPCs under replicative stress

We next examined whether exogenous addition of p16^Ink4a^ could recover the defects caused by *Jmjd3* deficiency. We used p16^INK4a^ HIV trans-activating protein (p16-TAT) that enters cells at high efficiency (Krosl *et al*, 2003). We first confirmed that p16-TAT was detectable in cultured cells for at least 12 h (Fig 6A) following its addition into the conditioned medium. We then allowed *Jmjd3^+/+^* or *Jmjd3^Δ/Δ^* LK cells treated with BSA or p16-TAT to form colonies and conducted BMT assays (Fig 6B). Addition of p16-TAT successfully restored the impaired colony forming ability of BSA treated *Jmjd3^Δ/Δ^* cells (Fig 6C). Furthermore, p16-TAT treated *Jmjd3^Δ/Δ^* cells partially rescued donor-derived chimeras in PB without affecting lineage differentiation of BMT recipients (Fig 6D and 6E), and almost fully rescued donor-derived chimerism in various BM cell fractions (Fig 6F). Moreover, the Lin^−^ population percentage in p16-TAT treated *Jmjd3^Δ/Δ^* cells in the BM was comparable to BSA treated *Jmjd3^+/+^* cells (Fig 6G).

**Figure 6.**
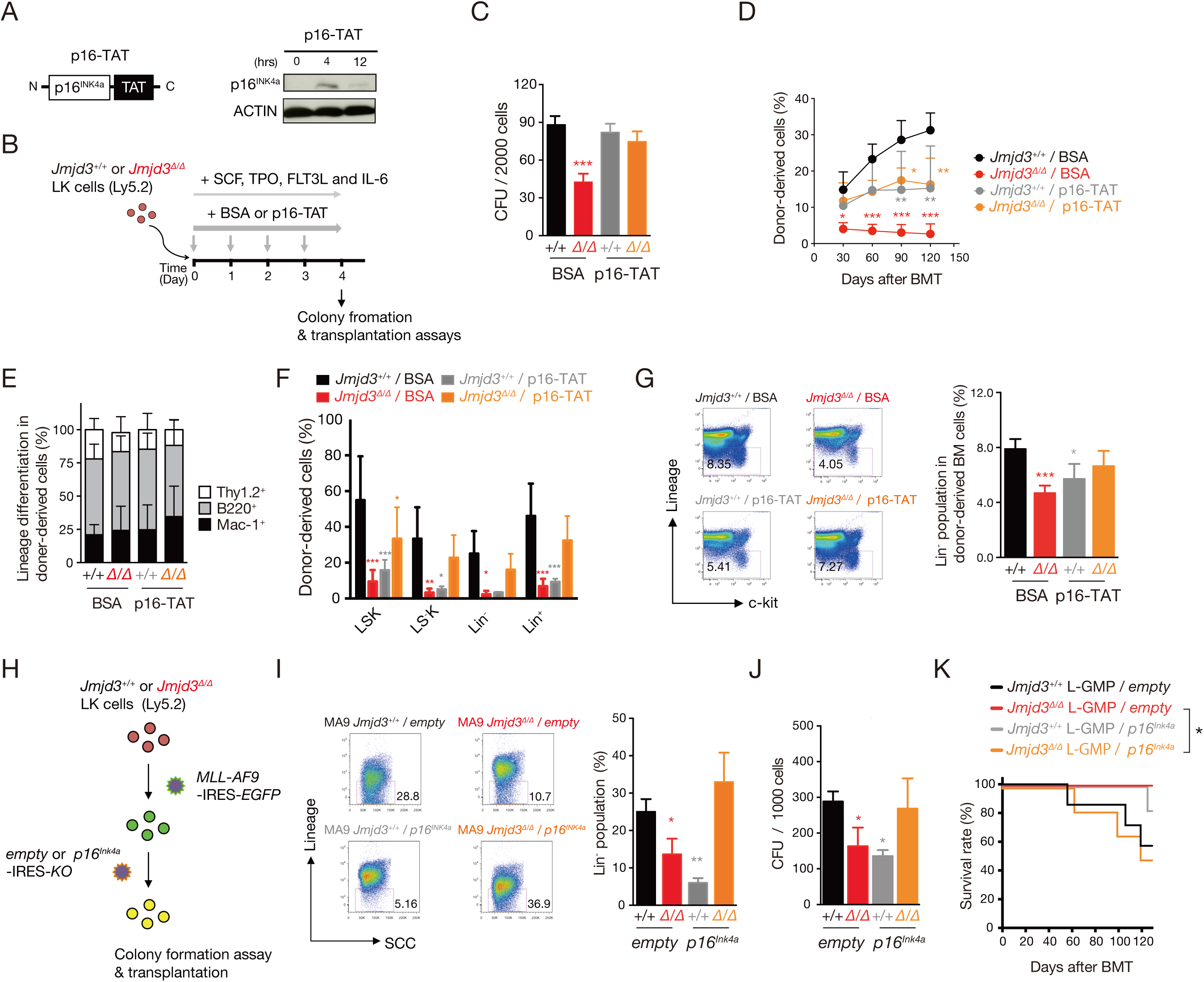
Rescue of *Jmjd3^Δ/Δ^*HSPCs stress-induced defects by exogenous and retroviral addition of *p16^Ink4a^*. (A) Schematic diagram of the p16^INK4a^-TAT fusion protein (p16-TAT, left panel) and immunoblot showing time-dependent stability of p16-TAT (50 nM) in LK cells. (B) Schematic of experimental procedure. *Jmjd3^+/+^* and *Jmjd3^Δ/Δ^* LK cells treated with BSA or p16-TAT (50nM) in a cytokine cocktail were subjected to colony forming and BMT assays. (C) Colony formation assay of *Jmjd3^+/+^* and *Jmjd3^Δ/Δ^* LK cells treated with BSA or p16-TAT for 4 days (mean + SD, n =3). (D) Donor-derived chimerism in the PB of recipient mice. 5.0×10^4^ *Jmjd3^+/+^* or *Jmjd3^Δ/Δ^* LK cells treated with BSA or p16-TAT for 4 days were transplanted into lethally irradiated recipients with 2.5×10^5^ wild type competitor MNBM cells for radioprotection (mean + SD, n = 4). (E) Percent of donor-derived, lineage committed (Thy1.2^+^, B220^+^, or Mac-1^+^) cells in the PB of recipient mice 4 months after BMT (mean + SD, n = 4). (F) Percent of donor-derived LSK, LS^−^K, Lin^−^, and Lin^+^ cells in the BM of recipient mice at 4 months after BMT (mean + SD, n = 4). (G) Flow cytometric profiles and percentages of Lin^−^ cells in the donor-derived BM of recipient mice 4 months after BMT (mean + SD, n = 4). (H) Schematic diagram of *MLL-AF9* and *p16^Ink4a^* co-transduction. LK cells from *Jmjd3^+/+^*and *Jmjd3^Δ/Δ^* mice were transfected with the *MLL-AF9-*IRES*-EGFP* retrovirus. EGFP^+^ cells were further transfected with *empty-* or *p16^Ink4a^*-IRES*-KO* (*Kusabira Orange*) retrovirus, and double-positive cells were subjected to the following assays. (I) Flow cytometric profiles of Lin^−^ cells in *MA9 Jmjd3^+/+^* and *Jmjd3^Δ/Δ^* leukemic cells transfected with *empty* or *p16^Ink4a^* (mean + SD, n = 3). (J) Colony forming assay of *Jmjd3^+/+^* and *Jmjd3^Δ/Δ^* L-GMPs transfected with *empty* or *p16^Ink4a^*. Bars indicate the colony numbers at the third round of replating (mean + SD, n = 3). (K) Kaplan–Meier survival curves of mice transplanted with *Jmjd3^+/+^* and *Jmjd3^Δ/Δ^* L-GMPs transfected with *empty* or *p16^Ink4a^*. 2.0×10^2^ L-GMPs were transplanted into lethally irradiated recipients with 2.5×10^5^ wild type competitor MNBM cells (n = 6–7). \**p* < 0.05, \*\**p* < 0.01, \*\*\**p* < 0.001.

p16^INK4a^ inhibits CyclinD1-CDK4/CDK6 complex formation (Ewen *et al*, 1994). We then examined cell cycle inhibition in *Jmjd3^Δ/Δ^* cells using a dominant negative form of CyclinD1, CyclinD1^T156A^, that is catalytically inactive and unable to phosphorylate RB (Diehl *et al*, 1997). *Empty*, *Ccnd1^WT^* or *Ccnd1^T156A^* coupled with *EGFP*, was introduced into *Jmjd3^+/+^* or *Jmjd3^Δ/Δ^* LK cells, and EGFP^+^ cells were transplanted into recipient mice (Fig EV4A). Intriguingly, although there was less chimerism in *Ccnd1^WT^*transduced *Jmjd3^Δ/Δ^* cells than in *empty* transduced *Jmjd3^Δ/Δ^* cells, *Ccnd1^T156A^* transduced *Jmjd3^Δ/Δ^*cells exhibited nearly the same repopulation and differentiation abilities as *empty* transduced *Jmjd3^+/+^* cells (Fig EV4B and EV4C). This finding was observed despite incomplete rescue of donor-derived percentages in BM subfractions (Fig EV4D). These results collectively indicate that introduction of cellular senescence by exogenous addition of p16^INK4a^ or retroviral transduction of an inactive form of CyclinD1 is capable of restoring and maintaining the stem cell potential of *Jmjd3^Δ/Δ^* HSPCs under replicative stress.

We next attempted to rescue impaired LSC potential due to *Jmjd3* deficiency using retroviral transduction of *p16^Ink4a^*. *Jmjd3^+/+^* and *Jmjd3^Δ/Δ^* LK cells were transduced with *MLL-AF9* coupled with *EGFP*. EGFP^+^ cells were transduced with *empty* or *p16^Ink4a^* coupled with *Kusabira Orange* (*KO*), and double-positive cells were subjected to colony replating and BMT assays (Fig 6H). The reduced Lin^−^ population in *MLL-AF9*-expressing *Jmjd3^Δ/Δ^* cells was successfully rescued by *p16^Ink4a^* transduction (Fig 6I). In addition, colonies of *Jmjd3^Δ/Δ^* L-GMPs were restored by the introduction of *p16^Ink4a^* (Fig 6J). Moreover, *p16^Ink4a^*expressing *Jmjd3^Δ/Δ^* L-GMPs exhibited comparable *in vivo* leukemogenic potential relative to *Jmjd3^+/+^* L-GMPs, as evidenced by similarly shortened survival rates of recipient mice (Fig 6K). Taken together, retroviral transduction of *p16^Ink4a^* successfully rescued the impaired leukemic potential of *MLL-AF9*-transformed *Jmjd3^Δ/Δ^*HSPCs via cellular senescence.

### Roles of JMJD3 in aging related accumulation of *p16^Ink4a^*

Accumulating evidence links senescence and aging to epigenetic alterations (Sen *et al*, 2016), and p16^INK4a^ is reported to be closely associated with stem cell aging (Janzen *et al*, 2006; Krishnamurthy *et al*, 2006; Molofsky *et al*, 2006). We first examined the expression of *p16^Ink4a^* in young and aged *Jmjd3^+/+^* and *Jmjd3^Δ/Δ^* HSPCs (Fig 7A). We observed aging associated upregulation of *p16^Ink4a^*in *Jmjd3^+/+^* HSPCs (Fig 7B), as previously reported (Janzen *et al*, 2006). Interestingly, no such effects were detected in *Jmjd3^Δ/Δ^* cells. In addition, significant enrichment of H3K27me3 was detected at the *p16^Ink4a^*promoter in aged *Jmjd3^Δ/Δ^* HSPCs compared with aged *Jmjd3^+/+^*cells (Fig 7C). We then performed competitive repopulation assays using LT-HSCs of young (2M), mature (18M) and aged (26M) *Jmjd3^+/+^* and *Jmjd3^Δ/Δ^*mice. When compared with *Jmjd3^+/+^* cells, the PB chimerism of *Jmjd3^Δ/Δ^* LT-HSCs was significantly lower at 2M, was comparable at 18M, and was significantly increased in early phases following BMT at 26M (Fig 7D), closely resembling the results reported in *p16^Ink4a^* deficient HSCs (Janzen *et al*, 2006). These findings indicate that *Jmjd3* contributes not only to JMJD3-p16^INK4a^ axis mediated cellular senescence, but also aging associated accumulation of *p16^Ink4a^* in a demethylase dependent manner that is overridden upon *Jmjd3* deficiency.

**Figure 7.**
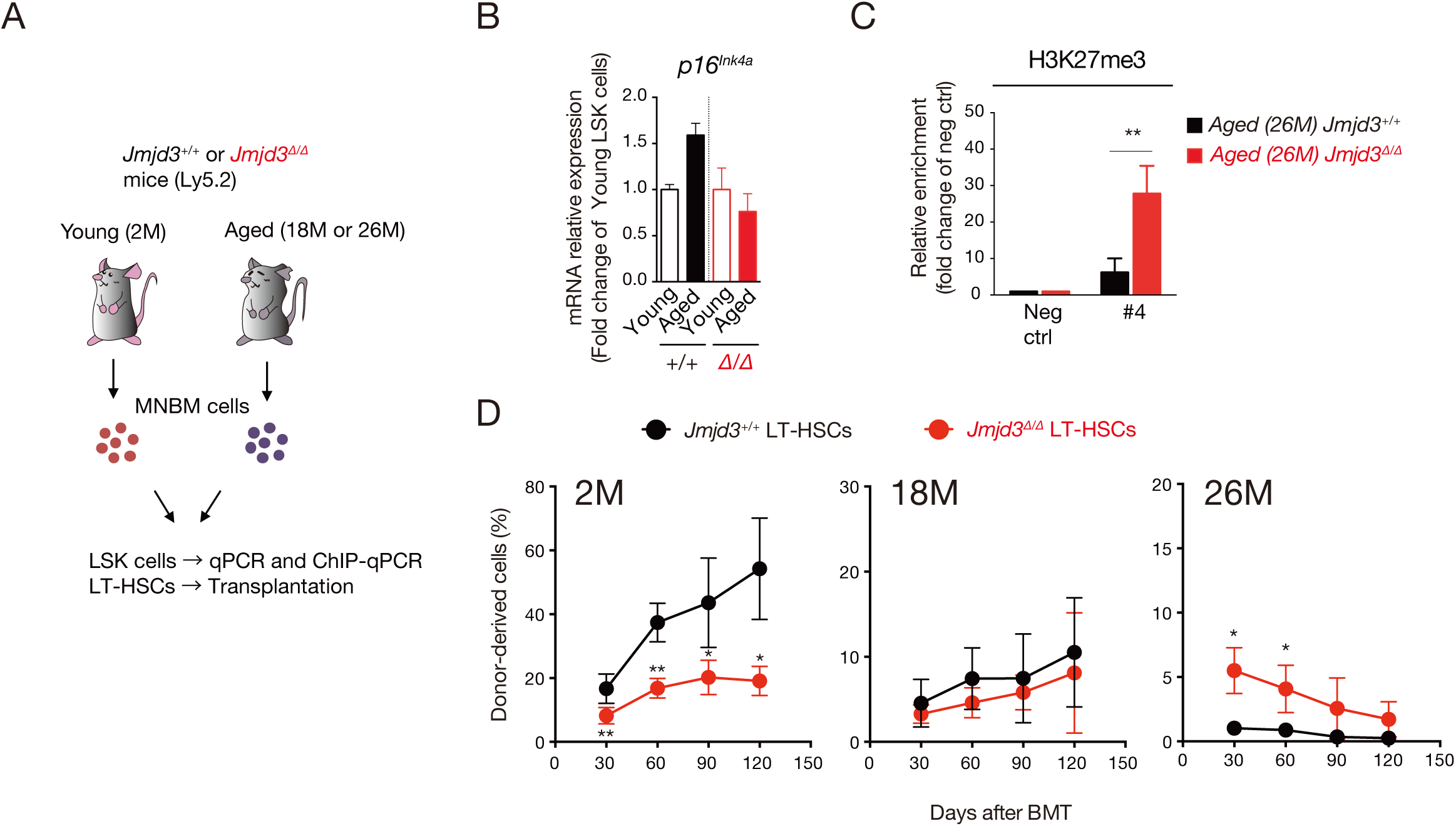
JMJD3 contributes to the aging process by regulating *p16^Ink4a^*. (A) Schematic diagram of the experiments with young and aged LSK cells and LT-HSCs. (B) qPCR analysis of *p16^Ink4a^* in LSK cells of young (2M) and aged (26M) *Jmjd3^+/+^* and *Jmjd3^Δ/Δ^* mice (mean + SD, n = 3). (C) H3K27me3 enrichment in the promoter region of *p16^Ink4a^* (#4, see Fig 3D) in LSK cells from aged (26M) *Jmjd3^+/+^* and *Jmjd3^Δ/Δ^* mice. Results are shown as fold changes relative to a negative control (Neg ctrl) (mean + SD, n = 3). (D) Competitive repopulation assays. 5.0×10^2^ LT-HSCs from young (2M), mature (18M), and aged (26M) *Jmjd3^+/+^*and *Jmjd3^Δ/Δ^* mice were transplanted into lethally irradiated recipients with 2.5×10^5^ competitor MNBM cells (mean ± SD, n = 6–7). \**p* < 0.05, \*\**p* < 0.01.

### GSK-J4, a JMJD3 inhibitor, suppresses LSC potential by inhibiting JMJD3-p16^INK4a^ axis

Given that almost no changes were detected in steady state hematopoiesis by JMJD3 deficiency, we reasoned that functional inhibition of JMJD3 during cellular senescence may be effective against leukemogenesis and lead to attenuation of LSC potential without major effects on normal hematopoiesis. Thus, we hypothesized that GSK-J4, a JMJD3 demethylase inhibitor, is a potential treatment option for leukemia.

To explore whether GSK-J4 is a promising therapeutic drug for leukemogenesis during cellular senescence, we transduced *MLL-AF9* into wild type LK cells and treated with the inhibitor at the early phase of leukemogenesis (Fig 8A). *MA9* cells treated with 10μM of GSK-J4 exhibited a significantly reduced stem cell population (Fig 8B). We next investigated transcriptional changes in L-GMPs treated with GSK-J4. Consistent with *Jmjd3^Δ/Δ^*L-GMPs, the expression *Cdkn2a* in GSK-J4 treated L-GMPs was suppressed (Fig 8C). Although no significant difference of global H3K27me3 levels was detected, we observed significant enrichment of H3K27me3 at the *p16^Ink4a^* promoter in GSK-J4 treated L-GMPs (Fig 8D and 8E). Similarly, in GSEA, there were many commonly downregulated pathways between GSK-J4 treated and *Jmjd3^Δ/Δ^* L-GMPs (Fig 8F and Table EV2) and the previously identified representative gene sets significantly suppressed by *Jmjd3* deficiency in stress induced HSPCs (Fig 5) were also negatively enriched in GSK-J4 treated L-GMPs (Fig 8G). In addition, excessive cell cycle progression was observed in GSK-J4 treated L-GMPs (Fig 8H). These data indicate that a JMJD3 inhibitor suppresses cellular senescence via impaired activation of *Cdkn2a* genes, consistent with features observed in *Jmjd3^Δ/Δ^* L-GMPs.

**Figure 8.**
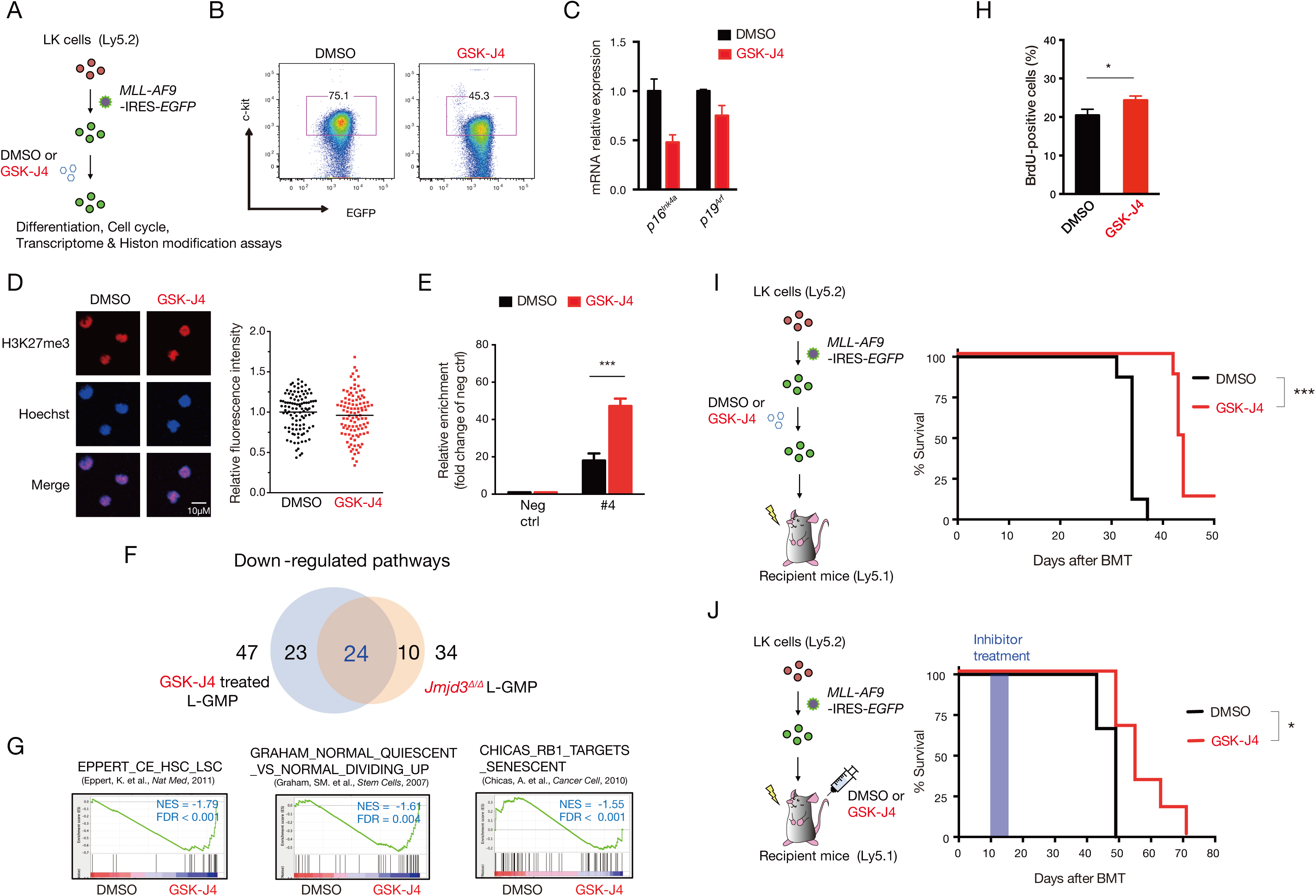
Inhibition of JMJD3 suppresses LSC potential via regulating *p16^Ink4a^* expression. (A) Schematic diagram of GSK-J4 treatment after *MLL-AF9* transduction. LK cells from wild type mice were transfected with the *MLL-AF9-*IRES*-EGFP* retrovirus. EGFP^+^ cells were further exposed to DMSO or GSK-J4 for 24h and subjected to the following assays. (B) Flow cytometric profiles of c-kit^+^ fractions in *MA9* cells treated with DMSO or GSK-J4 (5 or 10μM) for 24h. (C) qPCR analysis of *Cdkn2a* genes in L-GMPs exposed to DMSO or GSK-J4 (10μM) for 24h. (mean + SD, n = 3). (D) Immunofluorescence staining (left panel) and relative fluorescence intensity (right panel) of H3K27me3 in L-GMPs exposed to DMSO or GSK-J4 (10μM) for 24h. Mean values are indicated as bars (n = 105). (E) H3K27me3 levels in the promoter region of *p16^Ink4a^* (#4, see Fig 3D) in L-GMPs exposed to DMSO or GSK-J4 (10μM) for 24h. Results are shown as fold changes relative to a negative control (Neg ctrl) (mean + SD, n = 3). (F) Venn diagrams showing the overlap of negatively enriched KEGG pathways in L-GMPs exposed to GSK-J4 (10μM) for 24h and *Jmjd3^Δ/Δ^* L-GMPs. The overlapped pathways are listed in Table EV2. (G) GSEA plots of L-GMPs exposed to DMSO or GSK-J4 (10 μM) for 24h in the indicated gene sets (left, genes commonly upregulated in human HSC and LSC; middle, genes commonly upregulated in quiescent human CD34^+^ hematopoietic cells; right, genes commonly upregulated through the p16^INK4a^/RB1 pathway). Results are shown with NES and FDR values. (H) Flow cytometric analysis of BrdU incorporation in L-GMPs exposed to DMSO or GSK-J4 (5 or 10μM) for 24h. (mean + SD, n = 3). (I) Schematic diagram of *in vitro* GSK-J4 (10 μM) treatment for 24h after *MLL-AF9* transduction (left panel) and Kaplan–Meier survival plots of mice transplanted with these cells (right panel). 1.0×10^5^ *MA9* cells (Ly5.2^+^) were transplanted into lethally irradiated recipients with 2.5×10^5^ wild type competitor MNBM cells (n = 8). (J) Schematic diagram of *in vivo* GSK-J4 treatment after *MA9* transduction (left panel) and Kaplan–Meier survival plots of mice transplanted with *MA9* cells and treated *in vivo* (right panel). 1.0×10^5^ *MA9* cells (Ly5.2^+^) were transplanted into lethally irradiated recipients with 2.5×10^5^ wild type competitor MNBM cells. 10 days after BMT, DMSO or GSK-J4 (50 mg/kg/day) was intraperitoneally injected into the recipients for 5 consecutive days (n = 6).

Finally, we tested the effects of GSK-J4 on development of AML *in vivo*. The onset of AML was significantly delayed in recipient mice transplanted with *MA9* cells treated with GSK-J4 *in vitro* compared with those treated with DMSO (Fig 8I). Next, recipient mice transplanted with *MA9* cells were treated *in vivo* with DMSO or GSK-J4 by intraperitoneal injection at the early phase of leukemogenesis. We observed a significant increase in survival of treated mice (Fig 8J). This data indicates that selective inhibition of JMJD3 during cellular senescence can potentially suppress leukemic stem cell activity.

## Discussion

Cellular senescence is an intricate biological process with bilateral characteristics (Huang *et al*, 2022). The precise mechanisms behind cellular senescence in HSPCs and whether it positively or negatively controls stem cell functions are largely unknown. In this study, we highlighted the benefits of JMJD3 mediated cellular senescence in HSPCs, such as enhancing HSPC functional integrity to overcome stress induced hematopoietic defects through reduction of H3K27me3 at the *Cdkn2a* locus in a demethylase dependent manner, thereby upregulating the expression of *p16^Ink4a^*. JMJD3 mediated cellular senescence is considered a temporal and reversible cell cycle arrest. Recent works also point to cellular senescence as a critical step in promoting stem or oncogenic properties via a reversible cell cycle arrest and gene reprogramming under cellular senescence (Milanovic *et al*, 2018; Guccini *et al*, 2021).

The roles of the *Cdkn2a* in HSCs and LSCs remain controversial. Studies have shown that derepressed expression of *Cdkn2a* exerts deleterious effects on stem cell activity. HSCs and LSCs deficient in epigenetic genes, such as *Bmi1* and *Moz*, exhibit impaired self-renewal activity due to derepression of *Cdkn2a*, and this defect is rescued by genetic ablation of *Cdkn2a* (Oguro *et al*, 2006; Perez-Campo *et al*, 2013). In addition, overexpression of CyclinD1-CDK4, the inhibitory targets of p16^INK4a^, confers a competitive advantage to HSCs (Mende *et al*, 2015). In contrast, other reports have demonstrated that p16^INK4a^ induced cell quiescence is essential for maintaining stem cell activity. A study showed that treatment of HSCs with a CDK4/6 inhibitor accelerates hematologic recovery by protecting HSCs from stress induced proliferative exhaustion (He *et al*, 2017). Another report demonstrated that *p16^INK4a^* expression is required for survival in human papillomavirus associated tumor cells (McLaughlin-Drubin *et al*, 2013).

During somatic reprogramming by the Yamanaka factors, p16^INK4a^ expression is also induced, however, removal of p16^INK4a^ high cells increases reprogramming efficiency (Grigorash *et al*, 2023), meaning that quantitative and temporal p16^INK4a^ expression is must be precisely regulated for its impact on stem cell integrity. In our data, during the early phases following BMT, including the first BMT and second replating after *MLL-AF9* transduction, *Jmjd3^Δ/Δ^* cells that insufficiently upregulated *Cdkn2a* demonstrated greater reconstitution and proliferative abilities (Fig 1B and 2B). In contrast, in late phases such as the second BMT and third replating after *MLL-AF9* transduction, *Jmjd3^Δ/Δ^*cells exhibited significantly reduced reconstitution and proliferative capacity (Fig 1C and 2B). These results collectively indicate that impaired derepression of *Cdkn2a* confers an initial growth advantage to HSCs and LSCs via accelerated cell cycle progression, but eventually impairs stem properties through over-proliferation, and finally results in functional depletion. Our findings demonstrate that the JMJD3-p16^INK4a^ axis plays essential roles in preventing excessive cell cycle progression and consequential exhaustion of HSCs under stress.

Studies have also shown that p16*^INK4a^* is involved in stem cell aging. Aging associated *p16^INK4a^* accumulation is observed in various types of tissue stem cells, including HSCs (Janzen *et al*, 2006; Krishnamurthy *et al*, 2006; Molofsky *et al*, 2006) (see also Fig 7), but the underlying mechanisms are poorly understood. We found that exogenous stresses upregulate *p16^Ink4a^* through JMJD3 mediated H3K27 demethylation. This observation suggests the possibility that incomplete p16^INK4a^ downregulation following stress, due to loss of PRC2 function, for instance, and repeated stresses such as replicative stress and DNA damage would gradually lead to p16 accumulation with age. This idea is supported by our finding that aged *Jmjd3^Δ/Δ^* HSCs does not exhibit accumulation of *p16^Ink4a^*, associated with increased H3K27me3 at the *p16^Ink4a^* promoter. Since *Jmjd3^Δ/Δ^*LT-HSCs behave similarly to *p16^Ink4a^* deficient cells in BMT assays, we hypothesize that JMJD3 functions as an upstream regulator of p16^INK4a^ in HSC aging.

Acute leukemia is relapsed in a majority of patients after chemotherapy and the relapse emerges from an immature, drug resistant population of cells, LSCs. The ultimate goal of leukemia treatment is to eradicate LSCs. By using an *MLL-AF9* induced leukemia model, we demonstrated that selective inhibition of JMJD3 under cellular senescence is promising for suppressing the maintenance and proliferative ability of LSCs through derepression of *p16^Ink4a^*in a demethylase activity dependent manner (Fig 8). Indeed, AML cells treated with chemotherapy are in an induced senescent-like phenotype and post-senescent cells give rise to relapsed AML with enhanced LSC activity (Duy *et al*, 2021). Thus, treatment with a JMJD3 inhibitor such as GSK-J4 alongside chemotherapy during the cellular senescence in leukemia cells may offer a promising therapeutic option. This concept also suggests that LSC activity is not fixed but rather flexible and modulated by epigenetic plasticity. Moreover, GSK-J4 has been used to treat not only leukemia but also solid cancers where it exerts anti-proliferative effects (Ntziachristos *et al*, 2014; Hashizume *et al*, 2014; Lochmann *et al*, 2018). Our results strongly provide experimental evidence that targeting JMJD3 elicits anti-cancer effects, at least, in part, through inhibiting cancer stem cell activity.

Methylation and demethylation of H3K27 are important for maintaining stem cell function and determination of proper cell fate. No obvious hematological changes were observed in *Jmjd3^Δ/Δ^* cells at steady state (Fig EV2), and *Jmjd3* deficiency induced limited alterations of global H3K27me3 and gene expression. It is controversial whether other epigenetic factors, such as UTX, another H3K27 demethylase, may support or compensate the function of JMJD3 in its absence to maintain normal hematopoiesis. To address this possibility, we compared RNA-seq data from *Jmjd3^Δ/Δ^* and *Utx^Δ/Δ^* LSK cells (Sera *et al*, 2021) and analyzed genes that were more than 2-fold up- or downregulated following *Jmjd3* or *Utx* deficiency. We found no substantial overlap in either upregulated or downregulated genes (Fig EV5), indicating that the target genes of JMJD3 and UTX are mutually exclusive and it is unlikely that the minimal changes following JMJD3 deficiency are due to compensatory effects by UTX.

In this study, we investigated the roles of JMJD3 in adult hematopoiesis and found that JMJD3-p16^INK4a^ axis mediated cellular senescence functions as a hematopoietic gatekeeper not only to protect HSPCs from excessive cell cycle entry and eventual exhaustion under stress, but also to enhance the stemness in HSPCs exposed to stress via its demethylase activity. On the other hand, JMJD3 has been shown to regulate HSPC self-renewal and leukemogenesis through the AP-1 transcription factor complex in a demethylase activity independent manner (Mallaney *et al*, 2019), suggesting that the demethylase independent activities of JMJD3 may have additional functions on the transcriptional network during cellular senescence. Our findings provide insights into the regulatory mechanisms of hematopoiesis through histone modifications and suggest that JMJD3 is a prospective target for stem cell aging research and anti-cancer stem cell therapies.

## Structured Methods

### Reagents and Tools Table

| Reagent/Resource | Reference or Source | Identifier or Catalog Number |
| --- | --- | --- |
| Experimental Models |  |  |
| *Jmjd3<sup>flox/flox</sup>* mice | This paper | N/A |
| *MxCre<sup>+</sup>* mice | Kühn R *et al*, 1995 | N/A |
| Recombinant DNA |  |  |
| *pMYs-MLL-AF9-IRES-EGFP* | Koide S *et al*, 2016 | N/A |
| *pMYs-mouse p16<sup>Ink4a</sup>-IRES-KO (Kusabira Orange)* | This paper | N/A |
| *pMYs-mouse Ccnd1<sup>WT</sup> - IRES-EGFP* | This Paper | N/A |
| *pMYs-mouse Ccnd1<sup>T156A</sup>- IRES-EGFP* | This paper | N/A |
| Oligonucleotides |  |  |
| Primers for qPCR | This Paper | Table EV3 |
| Primers for ChIP | This Paper | Table EV3 |
| Antibodies |  |  |
| Rat anti-mouse-B220-APC for FACS | BD Pharmingen | Cat#553092 |
| Rat anti-mouse-B220-Biotin for FACS | BD Pharmingen | Cat#559971 |
| Rat anti-mouse-B220- PE-Cy7 for FACS | BD Pharmingen | Cat#552772 |
| Anti-BrdU-APC for FACS | BD Pharmingen | Cat#552598 |
| Rat anti-mouse-c-kit-APC-H7 for FACS | BD Pharmingen | Cat#560185 |
| Anti-mouse-c-kit-beads fot MACS | Miltenyi Biotec | Cat#130-097-146 |
| Rat anti-mouse-CD135-PE for FACS | BD Pharmingen | Cat#553842 |
| Anti-mouse-CD16/32-BV510 for FACS | BD Pharmingen | Cat#740111 |
| Rat anti-mouse-CD16/32-PE for FACS | BD Pharmingen | Cat#553145 |
| Rat anti-mouse-CD34-BV421 for FACS | BD Pharmingen | Cat#562608 |
| Rat anti-mouse-CD34-eFluor660 for FACS | eBioscience | Cat#50-0341-82 |
| Rat anti-mouse-CD34-FITC for FACS | BD Pharmingen | Cat#553733 |
| Rat anti-mouse-CD4-Biotin for FACS | eBioscience | Cat#13-0042-82 |
| Rat anti-mouse-CD8a-Biotin for FACS | BD Pharmingen | Cat# 553029 |
| Anti-mouse-Gr-1-APC for FACS | Biolegend | Cat#108412 |
| Rat anti-mouse-Gr-1-Biotin for FACS | BD Pharmingen | Cat#559971 |
| Rat anti-mouse-Gr-1-PE-Cy7 for FACS | BD Pharmingen | Cat#565033 |
| Mouse anti-mouse-Ly5.1-APC for FACS | BD Pharmingen | Cat#558701 |
| Mouse anti-mouse-Ly5.2-PE for FACS | BD Pharmingen | Cat#560695 |
| Rat anti-mouse-Mac-1-Biotin for FACS | BD Pharmingen | Cat#559971 |
| Rat anti-mouse-Mac-1-FITC for FACS | BD Pharmingen | Cat#553310 |
| Rat anti-mouse-Sca-1-PE-Cy7 for FACS | BD Pharmingen | Cat#558162 |
| Anti-Streptavidin-PerCP-Cy5.5 for FACS | BD Pharmingen | Cat#551419 |
| Anti-Streptavidin-beads for MACS | Miltenyi Biotec | Cat#130-048-101 |
| Rat anti-mouse-Ter119-Biotin for FACS | BD Pharmingen | Cat#559971 |
| Rat Anti-mouse-Thy1.2-FITC for FACS | BD Pharmingen | Cat#553004 |
| Mouse Anti-β-ACTIN for WB | Sigma Aldrich | Cat#A5316 |
| Rabbit anti-BMI1 for ChIP | Bethyl | Cat# A301-694A |
| Rabbit anti-H3K27me3 for IF, ChIP and CUT&RUN | Cell Signaling | Cat#9733 |
| Rabbit anti-JMJD3 for ChIP | Abcam | Cat#ab85392 |
| Rabbit anti-JMJD3 for WB | Cell Signaling | Cat#3457 |
| Rabbit anti-p16<sup>INK4a</sup> for WB | Cell Signaling | Cat#4824 |
| Rabbit anti-IgG for CUT&RUN | Cell Signaling | Cat#66362 |
| Bacterial and virus strains |  |  |
| Stbl2 Competent cells | Thermo Fisher | Cat#10268019 |
| Chemicals, peptides, and recombinant proteins |  |  |
| GSK-J4 | MCE MedChemExpress | Cat#HY-15648B |
| Polyinosinic-polycytidylic acid (pIpC) | Sigma | Cat#P1530 |
| Hoechst 33342 | Dojindo Laboratories | Cat#346-07951 |
| BrdU | BD Biosciences | Cat#552598 |
| Adriamycin | Wako | Cat#040-21521 |
| Z-VAD-FMK | Selleck | Cat#S7023 |
| Recombinant murine SCF | Peprtech | Cat#250-03 |
| Recombinant murine IL-3 | Peprtech | Cat#213-13 |
| Recombinant murine IL-6 | Peprtech | Cat#216-16 |
| Recombinant murine GM-CSF | Peprtech | Cat#250-05 |
| Recombinant murine TPO | Peprtech | Cat#315-14 |
| Recombinant murine FLT3-L | Peprtech | Cat#250-31L |
| p16-TAT peptide | Prospec | Cat#PKA-337 |
| Critical commercial assays |  |  |
| ChIP-IT Express Enzymatic Kit | Active Motif | Cat#53009 |
| Cut&Run assay kit | Cell Signaling | Cat#86652 |
| NEBNext Ultra II DNA Library Prep Kit | NEB | Cat#E7370L |
| Senescence β-Galactosidase Staining Kit | Cell Signaling | Cat#9860 |
| TruSeq RNA Sample Preparation Kit v2 | Illumina | Cat#RS-122-2001 |
| SureSelect Strand-Specific mRNA Library Preparation kit | Agilent Technologies | Cat#G9691B |
| BrdU Flow Kit | BD Biosciences | Cat#552598 |
| SuperScript VILO master mix kit | Invitrogen | Cat#11755050 |
| Software |  |  |
| FlowJo software | BD Biosciences | https://www.flowjo.com/solutions/flowjo |
| ImageJ64 | National Institutes of Health | https://imagej.nih.gov/ij/index.html |
| GSEA | Broad Institute | https://www.gsea-msigdb.org/gsea/index.jsp |
| Genome Browser | University of California Santa Cruz | https://genome.ucsc.edu |
| Transcription Factor and Histone ChIP-Seq pipeline | ENCODE | https://github.com/ENCODE-DCC/chip-seq-pipeline2 |
| Homer v4.11 | University of California San Diego | http://homer.ucsd.edu/homer/ |
| GraphPad Prism version 6 | GraphPad | https://www.graphpad.com |

## Methods

### Construction of the targeting vector and generation of conditional knockout mice

A bacterial artificial chromosome clone containing the mouse *Jmjd3* gene was purchased from the BACPAC Resource Center, Children’s Hospital Oakland Research Institute (Oakland). Fragments from the Aor51HI site in intron 11 to the artificially introduced BamHI site in intron 14 and from the SnaBI site in intron 17 to the BamHI site in intron 20 were used as the 5′ and 3′ arms of the targeting vector, respectively. A fragment from the artificially introduced BamHI site in intron 14 to the SnaBI site in intron 17 that contains exons 15–17 was *floxed* and inserted between the two arms, together with an *Frt*-flanked *Neomycin* (*Neo*)*-*resistance gene. A *diphtheria toxin A* (*DTA*) gene was attached to the 5′ end of the vector as a negative selector. KY1.1 ES cells (kindly provided by Dr. Junji Takeda, Osaka University, Japan) were electroporated with the linearized vector. Individual clones were screened via 5′ Southern blot using a 5′ probe and 3′ genomic PCR using P1 and P2 primers. Correctly targeted ES cells were microinjected into the blastocysts derived from C57BL/6 × BDF1 mice, and the resultant chimeric male mice were crossed with C57BL/6 female mice to transmit the mutant allele to progeny. The *Neo* resistance gene was removed by crossing heterozygotes with CAG-FLPe transgenic mice (RIKEN BRC, RBRC1834). The resultant *Jmjd3^+^*^/*flox*^ mice were crossed to *Mx*-*1Cre* mice, and the activation of Cre was induced by an intraperitoneal injection of polyinosinic-polycytidylic acid (pIpC). Mice that were backcrossed to C57BL/6N for at least five generations were used for experiments. Mouse experiments were performed in strict accordance with the Guide for the Care and Use of Laboratory Animals of the Hiroshima University Animal Research Committee (permission No. 29-111) and the Institute for Laboratory Animals of Tokyo Women’s Medical University Animal Research Committee (permission No. AE23-065).

### Flow cytometric analysis

Mononuclear (MN) cells were isolated from bone marrow (BM) and peripheral blood (PB) cells. After incubation with Fc-blocker (BD Biosciences), MNCs were stained with antibodies The surface marker phenotypes of each hematopoietic fraction are summarized in Table S1. Flow cytometry was performed on a FACSCanto II and a FACSAria II, and data were analyzed with FlowJo software (BD Biosciences).

### Bone marrow transplantation (BMT) assay

*Jmjd3^+/+^* or *Jmjd3^Δ/Δ^* HSPCs (Ly5.2), together with competitor BM MN cells (Ly5.1) for radioprotection, were transplanted intravenously into recipient mice (Ly5.1) lethally irradiated at a dose of 8.0 Gy, as previously described (Nakata *et al*, 2017). The chimerism of donor-derived hematopoietic cells was monitored by flow cytometry.

### Western blot

Whole cell lysates were run on SDS-PAGE gels and transferred onto a PVDF membrane (Millipore). Non-specific binding was blocked with 5% skim milk in Tris-buffered saline with 0.1% Tween-20 (TBS-T), followed by an overnight reaction with primary antibodies. Immunoreactive proteins were identified with HRP-conjugated anti-rabbit IgG (GE Healthcare) and anti-mouse IgG (GE Healthcare), and reacted with chemiluminescent HRP substrate (Millipore).

### Retroviral transduction

LK cells were first stained by anti-lineage-biotin and enriched by MACS LS-columns, streptavidin microbeads, and anti-c-kit microbeads. For transduction of *MLL-AF9* and/or *p16^Ink4a^*, LK cells were cultured for 3 days in RPMI1640 medium containing 10% FCS supplemented with 20 ng/ml SCF, 10 ng/ml IL-3, and 10 ng/ml IL-6. The cells were infected with retrovirus carrying *MLL-AF9*-IRES*-EGFP* and/or *p16^Ink4a^*-IRES*-KO* (*Kusabira Orange*). For transduction of *Ccnd1*, LK cells were cultured for 3 days in S-clone SF-O3 serum-free medium (EIDIA Co) containing 1% BSA supplemented with 100 ng/ml SCF and 100 ng/ml TPO, and the cells were transduced with retrovirus carrying mouse *Ccnd1^WT^*- or mouse *Ccnd1^T156A^*-IRES*-EGFP*.

### Colony formation and serial replating assays

Cells were seeded in MethoCult^TM^ M3231 (STEMCELL Technologies), supplemented with cytokines (PeproTech). For normal HSPCs, 50 ng/ml SCF, 50 ng/ml TPO, and 50 ng/ml FLT3L were used. For *MLL-AF9*-transformed LK cells, 20 ng/ml SCF, 10 ng/ml IL-3, 10 ng/ml IL-6, and 10 ng/ml GM-CSF were used. Colony numbers were counted at 4–6 days after plating, and the cells were collected and replated.

### Immunofluorescent staining

Cells were plated on poly-L-lysine-coated glass slides (Matsunami Glass), fixed with 4% paraformaldehyde and permeabilized with 0.5% Triton X-100. Non-specific binding was blocked with Protein block serum-free (Dako) and cells were stained with the primary antibody. After washing with PBS, the cells were stained with Alexa Fluor 488- or 555- conjugated goat anti-rabbit IgG antibody (Invitrogen). Nuclei were counterstained with Hoechst 33342. Images were taken with an FV-1000 confocal microscope (Olympus). Cells were chosen randomly in multiple fields, and fluorescence intensities of individual cells (more than 110 cells per experiment) were quantified computationally using ImageJ64 software.

### Quantitative real-time PCR

Total cellular RNA was isolated with TRIzol reagent (Invitrogen). RNA was reverse-transcribed with a SuperScript VILO master mix kit (Invitrogen), according to the manufacturer’s protocol. Quantitative real-time PCR was performed with power SYBR Green PCR Master Mix (Thermo Fisher Scientific) on a StepOnePlus Real-Time PCR system (Applied Biosystems) running StepOne 2.3 Software (Applied Biosystems). The primer sequences are listed in Table S3 All data are presented relative to *Hprt*.

### Transcriptome analysis and data processing

RNA-Seq libraries were prepared with the TruSeq RNA Sample Preparation Kit v2 and SureSelect Strand-Specific mRNA Library Preparation kit. Transcriptome analysis was performed using a next-generation sequencer (GAIIx and HiSeq 2500; Illumina), according to the manufacturer’s instructions. The sequence tags (more than 3 × 10^7^ reads for each sample) were mapped onto the mouse genomic sequence (UCSC Genome Browser, version mm10) with ELAND for DRA004290 and TopHat for DRA008581, DRA005628 and DRA010428. Normalized gene expression was compared between *Jmjd3^Δ/Δ^* and *Jmjd3^+/+^* LSK cells at steady state, LSK cells after BMT, and L-GMPs, and between DMSO and GSK-J4-treated L-GMPs, and the results were analyzed with GSEA software. Gene sets with a false discovery rate (FDR) q-value < 0.25 were considered statistically significant.

### Analysis of cell cycle activity

BrdU incorporation was analyzed with a BrdU Flow Kit. *Jmjd3^+/+^* or *Jmjd3^Δ/Δ^* mice were intraperitoneally injected with 1 mg of BrdU at 8, 16, and 24h before analysis. Cultured LK cells transfected with *MLL-AF9* were treated with BrdU (10μM) in cultured medium 8h before analysis. Cells were stained with antibodies of cell surface markers and then fixed, permeabilized, and stained with an anti-BrdU antibody according to the manufacturer’s instructions.

### ChIP-qPCR assay

Chromatin was enzymatically shred with the ChIP-IT Express Enzymatic Kit and immunoprecipitated overnight at 4°C with Protein G magnetic beads and antibodies, as listed in Table S3 Subsequently, chromatin was eluted, reverse cross-linked, and treated with Proteinase K, according to the manufacturer’s instructions. ChIP-qPCR data are presented as relative enrichment levels normalized to the *p21* promoter region as a negative control (Neg ctrl). Primer sequences are listed in Table S3

### Cut&Run assay and analysis

Anti-H3K27me3 and anti-IgG were used for Cut&Run. This assay was performed with the Cut&Run assay kit, according to the manufacturer’s protocol. Libraries were generated using the NEBNext Ultra II DNA Library Prep Kit for Illumina and sequenced on an Illumina NovaSeq 6000. Paired-end fastq files were processed using the ENCODE Transcription Factor and Histone ChIP-Seq pipeline. Reads were trimmed using cutadapt v2.5. and aligned to the mm10 genome using Bowtie2 v2.3.4.3., and SAMtools v1.9 was used to convert the output file to BAM format. Duplicates were removed using Picard Tools v2.20.7. Peak calling was performed with MACS2 v2.2.4, and the peaks were compared with IgG peaks before subsequent analysis. Bigwig files were normalized by a scaling factor based on the read counts of spike-in DNA using Deeptools v3.3.1 bamCoverage tools and visualized in the UCSC genome browser. Homer v4.11 was used for peak annotation and motif analysis. Bedtools v2.29.0 intersect was used to determine peak overlaps and assign target genes.

### β-galactosidase staining assay

Senescence-associated β-galactosidase staining was performed using a Senescence β-Galactosidase Staining Kit. L-GMPs were cultured with 0.025μg/ml of adriamycin (ADR) and 10μM of Z-VAD-FMK (caspase inhibiter). The cells were washed with PBS, then fixed and stained with β-galactosidase substrate X-Gal according to the manufacturer’s instructions. The senescent cells were counted using ImageJ software.

### p16^INK4a^-TAT treatment

LK cells were cultured for 4 days in S-clone SF-O3 serum-free medium containing 1% BSA supplemented with cytokines (100 ng/ml SCF, 100 ng/ml TPO, 100 ng/ml FLT3L, and 50 ng/ml IL-6). BSA was then added to the cultured medium or 50nM of p16-TAT. Fresh BSA or p16-TAT (to 10% of total culture medium) was added every 24h. After incubation for 4 days, cells were subject to colony formation and BMT assays.

### GSK-J4 treatment

*MA9*-expressing cells were exposed to 5 or 10μM of GSK-J4 for 24h. After incubation for 24h, cells were subject to colony formation and BMT assays. For GSK-J4 treatment *in vivo*, *MA9*-expressing cells (Ly5.2) were transplanted intravenously into recipient mice (Ly5.1) lethally irradiated at a dose of 8.0 Gy with 2.5×10^5^ wild type competitor MNBM cells for radioprotection. 10 days after BMT, DMSO or GSK-J4 (50mg/kg/day) was intraperitoneally injected into the recipients for 5 consecutive days.

### Statistical analysis

Statistical differences between the means of two groups were assessed using two-tailed Student’s t-test. When multiple treatment groups were compared, statistical significance of differences was assessed using a one-way ANOVA followed by Dunnett’s test. Mouse survival curves were constructed using the Kaplan–Meier methodology and compared with a log-rank test. All statistical tests were performed with GraphPad Prism version 6.

## Acknowledgments

We thank Yuki Sakai, Sawako Ogata, Miho Koizumi and Megumi Nakamura for animal care, genotyping, and molecular experiments. This work was in part supported by Grant-in-Aid for Japan Society for the Promotion of Science Fellows, The Takeda Science Foundation, The Naito Foundation and AMED-CREST (JP22gm1310006).

## Author Contributions

**Yuichiro Nakata**: Conceptualization; supervision; data curation; investigation; funding acquisition; formal analysis; writing – review and editing. **Takeshi Ueda**: Conceptualization; supervision; data curation. **Yasuyuki Sera**: Formal analysis. **Akinori Kanai**: Data curation. **Ken-ichiro Ikeda**: Formal analysis. **Norimasa Yamasaki**: Resources. **Akiko Nagamachi**: Formal analysis. **Kohei Kobatake**: Formal analysis. **Masataka Taguchi**: Conceptualization. **Yusuke Sotomaru**: Resources. **Tatsuo Ichinohe**: Investigation. **Zen-ichiro Honda**: Investigation. **Ichiro Manabe**: Investigation. **Toshio Suda**: Investigation. **Keiyo Takubo**: Conceptualization; Methodology. **Osamu Kaminuma**: Conceptualization; supervision; resources. **Hiroaki Honda**: Conceptualization; supervision; resources; project administration; funding acquisition; writing – review and editing.

## Data Availability

All sequencing data has been deposited in the DNA Data Bank of Japan.

RNA-seq; DRA004290 (https://ddbj.nig.ac.jp/resource/sra-submission/ DRA004290), DRA008581 (https://ddbj.nig.ac.jp/resource/sra-submission/DRA008581), DRA005628 (https://ddbj.nig.ac.jp/resource/sra-submission/DRA005628) and DRA010428 (https://ddbj.nig.ac.jp/resource/sra-submission/DRA010428).

Cut & Run-seq; DRA016982 (https://ddbj.nig.ac.jp/resource/sra-submission/DRA016982).

---

**Figure EV1.**
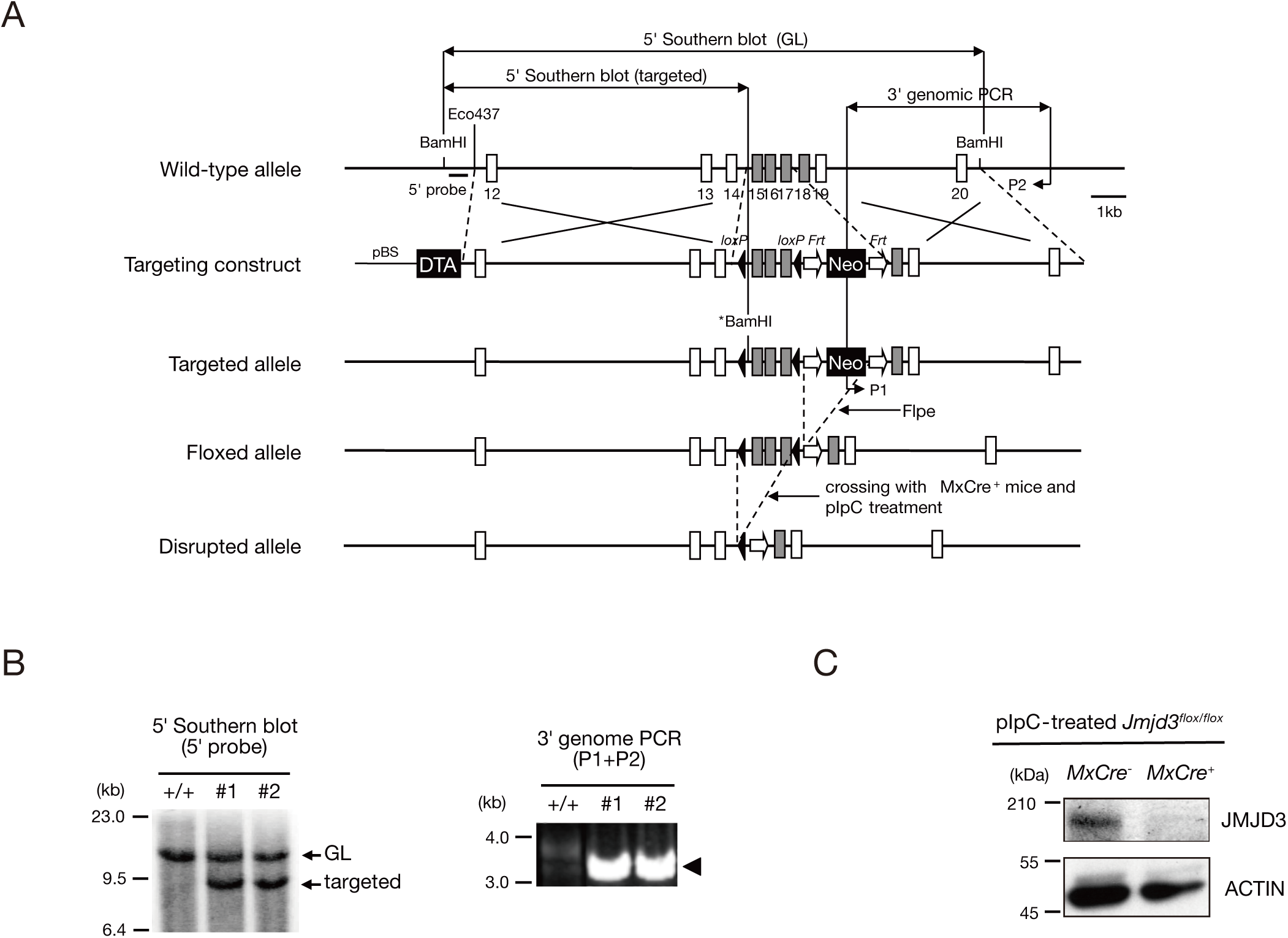
Targeting strategy, genotyping of ES clones, and deletion of *Jmjd3*. (A) Exons 15–17 of the mouse *Jmjd3* gene were encompassed by two *loxP* sites (black triangles), and a *neomycin*-resistance gene (*Neo*) was flanked by two *Frt* sites (white arrows). After removing *Neo* by Flpe, *floxed* exons were deleted by crossing with *MxCre*^+^ mice and pIpC treatment. The position of the genomic probe for the 5′ Southern blot (5′ probe), primers for the 3′ genomic PCR (P1 and P2), and the positions of restriction enzymes (BamHI and Eco437) are shown. BamHI with an asterisk (*BamHI) is an artificial enzyme site introduced by *in vitro* mutagenesis. Gray boxes indicate the exons that encode the JmjC domain. (B) Homologously recombined ES clones (#1 and #2) identified by 5′ Southern blot and 3′ genomic PCR. Germline (GL) and targeted bands in 5′ Southern blot are indicated by arrows (left panel) and PCR products for the 3′ genomic PCR are indicated by arrowheads (right panel). (C) Deletion of JMJD3 protein in BM cells of pIpC treated *Jmjd3^flox/flox^*;*MxCre*^+^ mice.

**Figure EV2.**
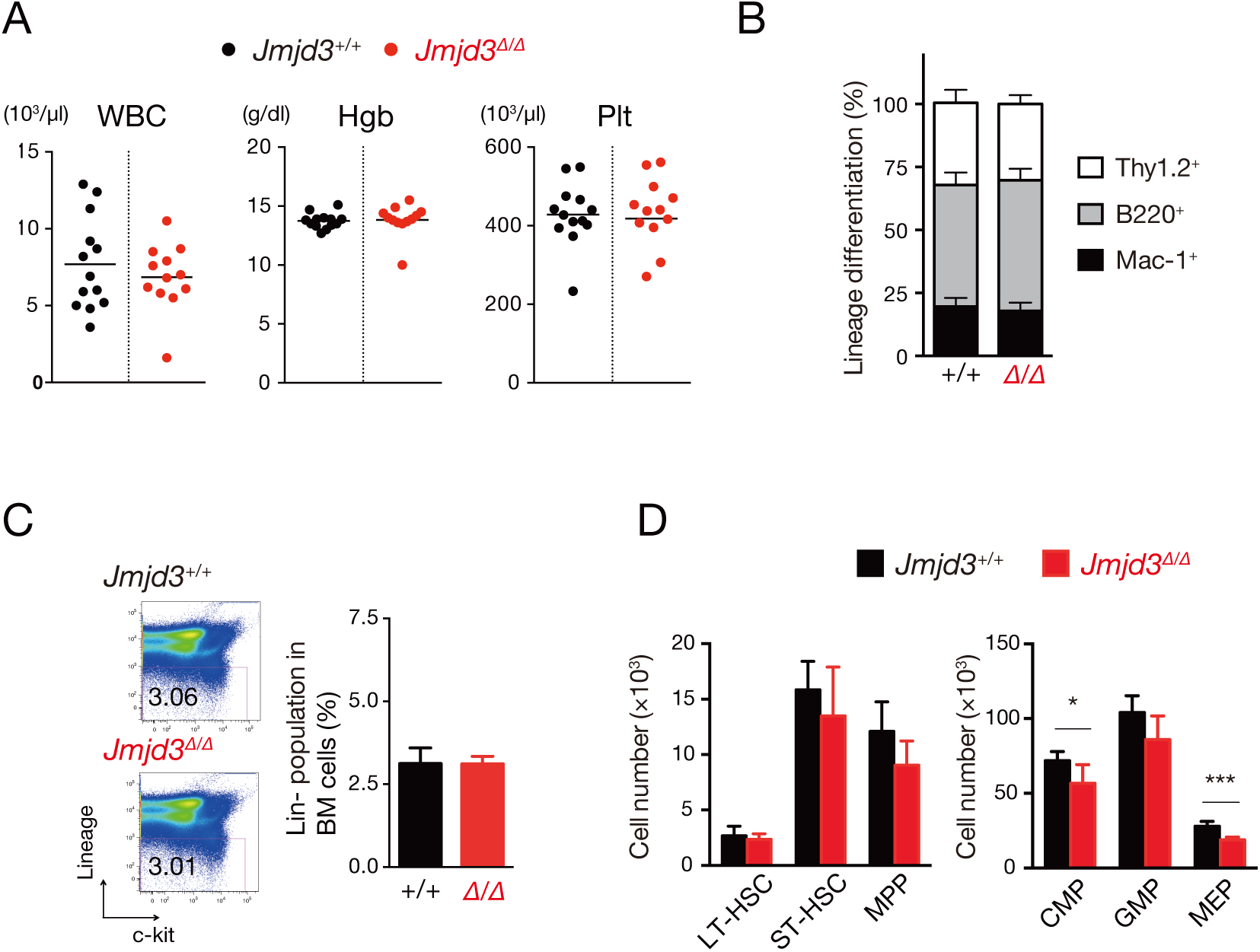
Analysis of *Jmjd3^Δ/Δ^* hematopoietic cells at steady state. (A) Analysis of PB parameters in *Jmjd3^+/+^* and *Jmjd3^Δ/Δ^*mice at 4 weeks after pIpC treatment. White blood cell (WBC) counts, hemoglobin concentration (Hgb), and platelet (Plt) number in the PB of *Jmjd3^+/+^* (n = 13) and *Jmjd3^Δ/Δ^* mice (n = 12) are plotted as dots, and the mean values are indicated as bars. (B) Analysis of lineage differentiation (Thy1.2^+^, B220^+^, and Mac-1^+^ cells) in the PB cells of *Jmjd3^+/+^* (n = 13) and *Jmjd3^Δ/Δ^* mice (n = 12) (mean ± SD). (C) Flow cytometric profiles of lineage^-^ (Lin^−^) cells in the BM of *Jmjd3^+/+^* and *Jmjd3^Δ/Δ^* mice (mean + SD, n = 5). (D) Absolute numbers of HSC subpopulations (LT-HSC, ST-HSC, and MPP) and myeloid progenitors (CMP, GMP, and MEP) in the BM of *Jmjd3^+/+^* and *Jmjd3^Δ/Δ^* mice (mean + SD, n = 5). \**p* < 0.05, \*\*\**p* < 0.001.

**Figure EV3.**
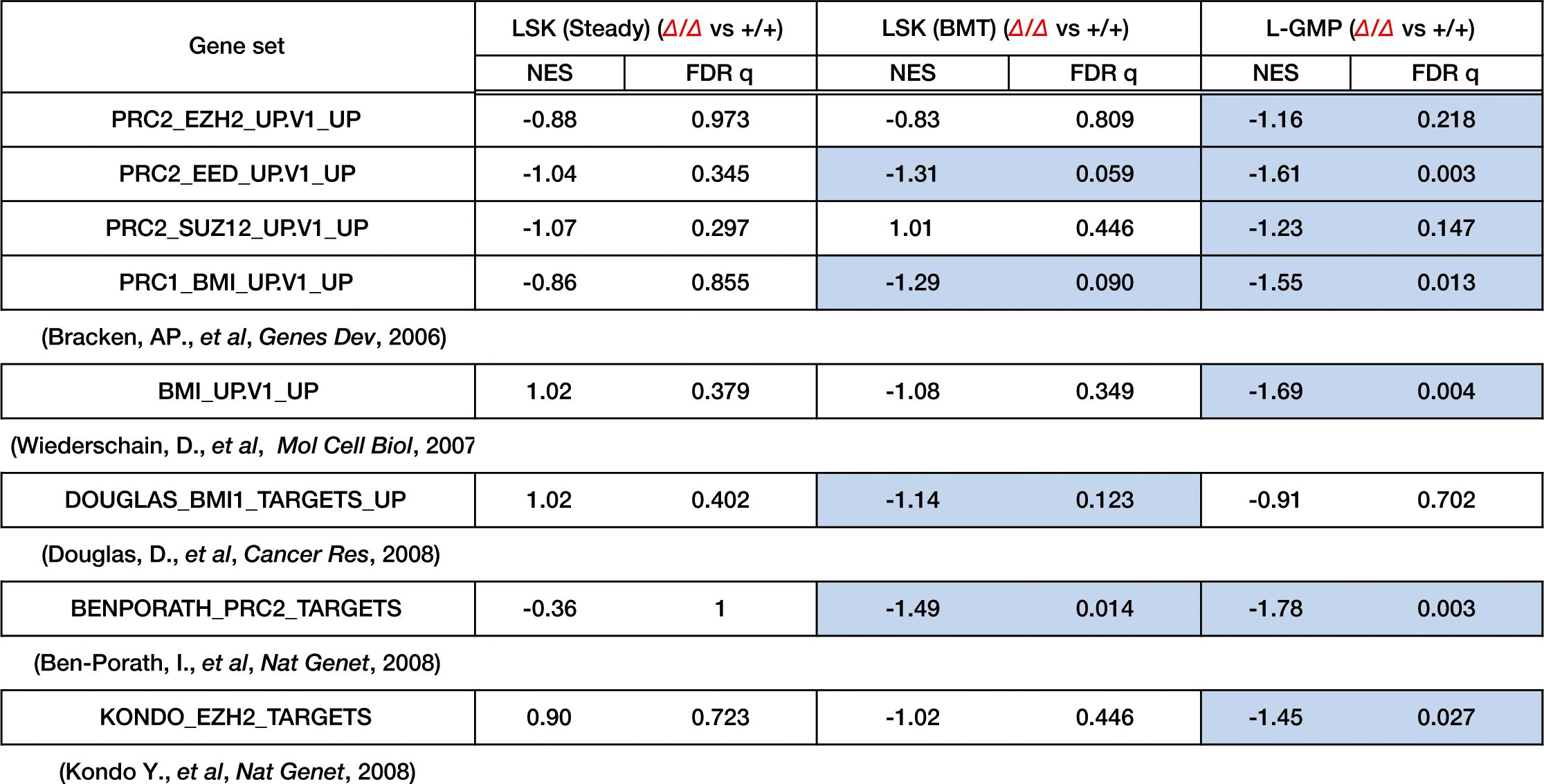
JMJD3 competitively regulates Polycomb targets under stress. Comparison of gene set enrichment between Polycomb proteins and JMJD3. Gene sets upregulated in cells deficient in the indicated Polycomb genes were compared with those downregulated in *Jmjd3*-deficient LSK (Steady), LSK (BMT), or L-GMP. NES and FDR are indicated. Blue boxes show significantly enriched pathways (FDR < 0.25).

**Figure EV4.**
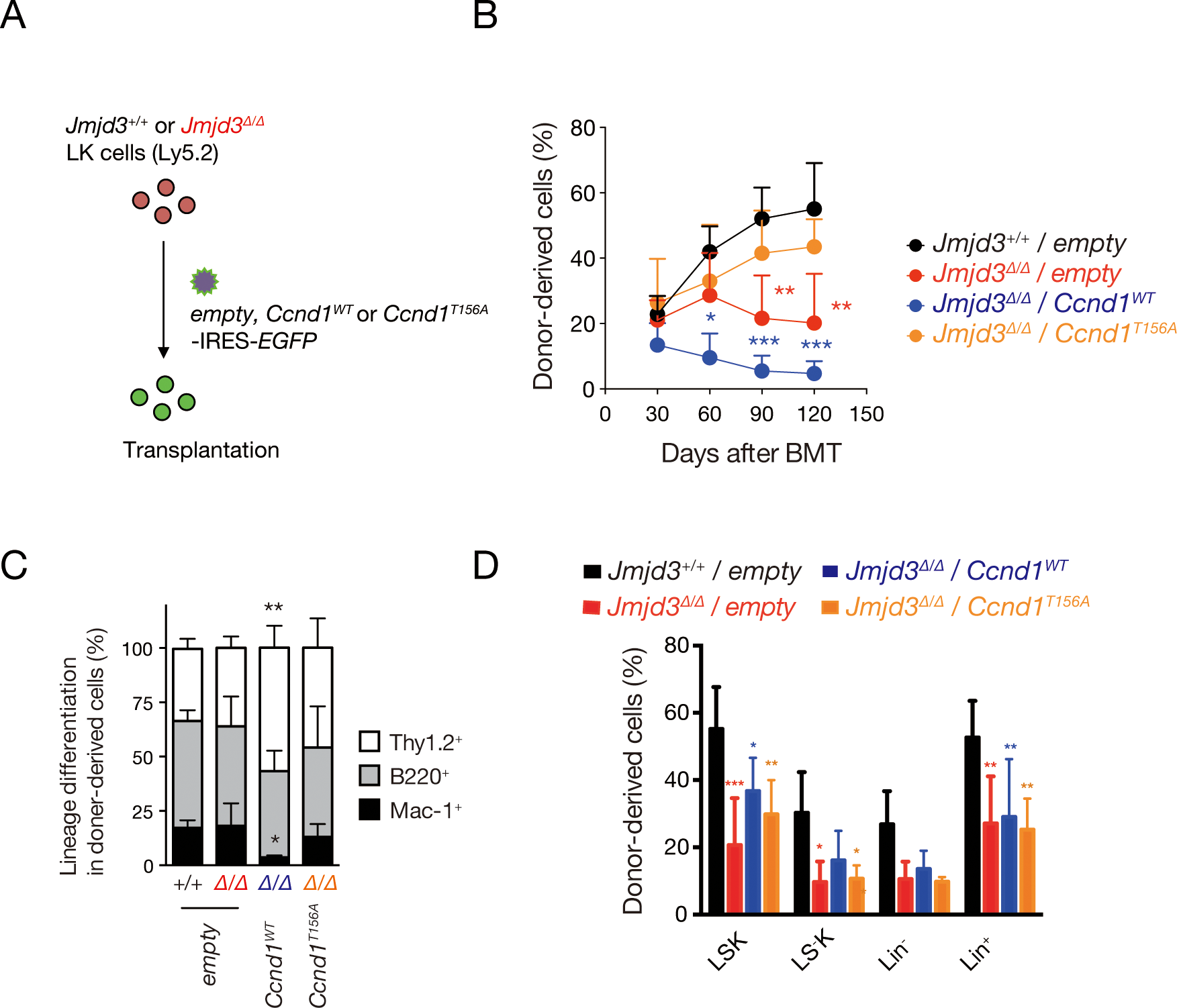
HSPC defects caused by *Jmjd3* loss under replicative stress were rescued by retroviral expression of a *Ccnd1* mutant. (A) A schematic diagram of *empty*, *Ccnd1^WT^*, and *Ccnd1^T156A^* transduction. LK cells from *Jmjd3^+/+^* and *Jmjd3^Δ/Δ^*mice were transfected with *empty*-, *Ccnd1^WT^*-, or *Ccnd1^T156A^*-IRES*-EGFP* retrovirus, and 5.0×10^4^ EGFP^+^ cells were transplanted into lethally irradiated recipients with 2.5×10^5^ wild type competitor MNBM cells for radioprotection (mean + SD, n = 4). (B) Chimerism of donor-derived PB cells in the recipients. Results of mice transplanted with *Jmjd3^+/+^*/*empty*, *Jmjd3^Δ/Δ^*/*empty*, *Jmjd3^Δ/Δ^*/*Ccnd1^WT^*, and *Jmjd3^Δ/Δ^*/*Ccnd1^T156A^* cells are shown (mean ± SD, n = 4). (C) Lineage differentiation in the donor derived PB cells in recipients 4 months after BMT (mean + SD, n = 4). (D) Chimerism of donor derived BM cells in recipients 4 months after BMT (mean ± SD, n = 4). \**p* < 0.05, \*\**p* < 0.01, \*\*\**p* < 0.001.

**Figure EV5.**
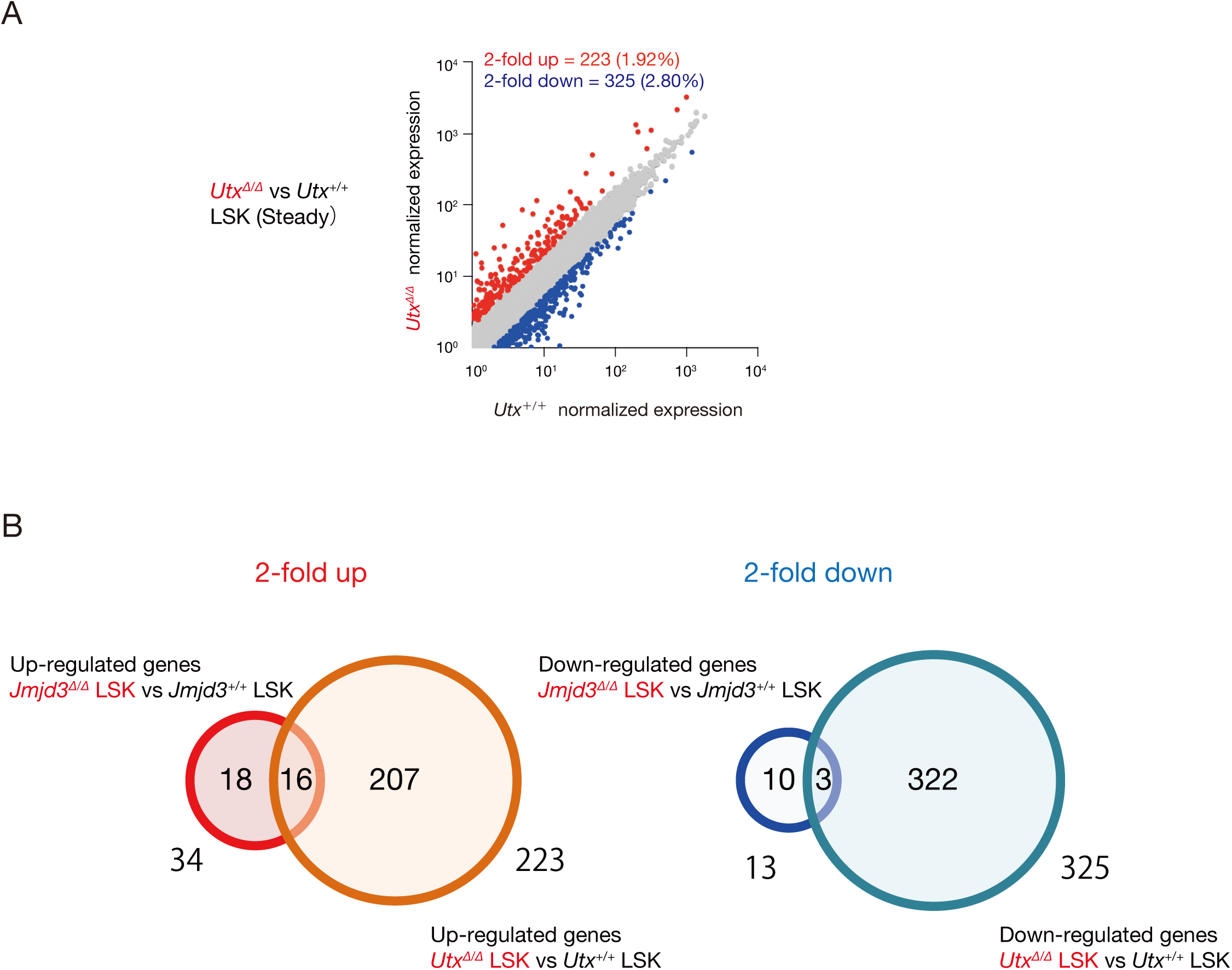
Overlap of genes up-regulated or down-regulated in *Jmjd3^Δ/Δ^* and *Utx^Δ/Δ^* LSK cells at steady state. (A) Scatter plots comparing normalized expression of individual genes (RPKM > 1) in LSK cells of *Utx^Δ/Δ^* mice compared with *Utx^+/+^* mice at steady state. Genes more than 2- fold up- and downregulated are plotted as red and blue dots, respectively. (B) Venn diagrams showing the overlap of genes more than 2-fold up- or downregulated in *Jmjd3^Δ/Δ^* and *Utx^Δ/Δ^*LSK cells at steady state.

**Table EV1.** Surface marker phenotypes to separate mouse hematopoietic stem/progenitor cells and leukemic stem cells.

| **Population** | **Surface phenotype** |
| --- | --- |
| CMP | CD34<sup>+</sup> CD16/32<sup>-</sup> c-kit<sup>+</sup> Sca-1<sup>-</sup> Lineage<sup>-</sup> |
| GMP and L-GMP | CD34<sup>+</sup> CD16/32<sup>+</sup> c-kit<sup>+</sup> Sca-1<sup>-</sup> Lineage<sup>-</sup> |
| LT-HSC | CD34<sup>-</sup> CD135<sup>-</sup> c-kit<sup>+</sup> Sca-1<sup>+</sup> Lineage<sup>-</sup> |
| MEP | CD34<sup>-</sup> CD16/32<sup>-</sup> c-kit<sup>+</sup> Sca-1<sup>-</sup> Lineage<sup>-</sup> |
| MPP | CD34<sup>+</sup> CD135<sup>+</sup> c-kit<sup>+</sup> Sca-1<sup>+</sup> Lineage<sup>-</sup> |
| ST-HSC | CD34<sup>+</sup> CD135<sup>-</sup> c-kit<sup>+</sup> Sca-1<sup>+</sup> Lineage<sup>-</sup> |

CMP, common myeloid progenitor; GMP, granulocyte/macrophage progenitor; L-GMP, leukemic GMP; LT-HSC, long-term HSC; MEP, Megakaryocyte/erythrocyte progenitor; MPP, multipotent progenitor; ST-HSC, short-term HSC.
CD4, CD8, B220, Mac1, Gr1 and Ter119 were used to exclude lineage committed cells.

**Table EV2.** Gene set enrichment analysis.

| Common negative enriched gene sets in GSK-treated and *Jmjd3*<sup>Δ/Δ</sup> L-GMP cells |  |
| --- | --- |
|  | Gene sets |
| 1 | KEGG_NEUROACTIVE_LIGAND_RECEPTOR_INTERACTION |
| 2 | KEGG_NATURAL_KILLER_CELL_MEDIATED_CYTOTOXICITY |
| 3 | KEGG_CALCIIUM_SIGNALING_PATHWAY |
| 4 | KEGG_FC_GAMMA_R_MEDIATED_PHAGOCYTOSIS |
| 5 | KEGG_LEUKOCYTE_TRANSENDOTHELIAL_MIGRATION |
| 6 | KEGG_JAK_STAT_SIGNALING_PATHWAY |
| 7 | KEGG_PHOSPHATIDYLINOSITOL_SIGNALING_SYSTEM |
| 8 | KEGG_FC_EPSILON_RI_SIGNALING_PATHWAY |
| 9 | KEGG_REGULATION_OF_ACTIN_CYTOSKELETON |
| 10 | KEGG_FOCAL_ADHESION |
| 11 | KEGG_ARRHYTHMOGENIC_RIGHT_VENTRICULAR_CARDIOMYOPATHY_ARVC |
| 12 | KEGG_VASCULAR_SMOOTH_MUSCLE_CONTRACTION |
| 13 | KEGG_LONG_TERM_POTENTIATION |
| 14 | KEGG_MELANOGENESIS |
| 15 | KEGG_AXON_GUIDANCE |
| 16 | KEGG_T_CELL_RECEPTOR_SIGNALING_PATHWAY |
| 17 | KEGG_ADHERENS_JUNCTION |
| 18 | KEGG_PANCREATIC_CANCER |
| 19 | KEGG_B_CELL_RECEPTOR_SIGNALING_PATHWAY |
| 20 | KEGG_HEMATOPOIETIC_CELL_LINEAGE |
| 21 | KEGG_LONG_TERM_DEPRESSION |
| 22 | KEGG_ECM_RECEPTOR_INTERACTION |
| 23 | KEGG_TIGHT_JUNCTION |
| 24 | KEGG_NON_SMALL_CELL_LUNG_CANCER |

